# Structural and genetic dissection of RNA-guided gene repression by TnpB-derived transcription factors

**DOI:** 10.64898/2026.09.22.753285

**Authors:** Renjian Xiao, Americo A. Casas-Ciniglio, Hoang C. Le, Tanner Wiegand, Dan Xie, Leifu Chang, Samuel H. Sternberg

**Affiliations:** Department of Biological Sciences, Purdue University, 915 W. State Street, West Lafayette, IN 47907, USA; Department of Genetics and Development, Columbia University, New York, NY, USA; Department of Biochemistry and Molecular Biophysics, Columbia University, New York, NY, USA; Howard Hughes Medical Institute, Columbia University, New York, NY, USA; Purdue Institute for Cancer Research, Purdue University, West Lafayette, IN 47907, USA

## Abstract

TnpB nucleases, the evolutionary progenitors of CRISPR-associated Cas12 enzymes, are transposon-encoded, RNA-guided endonucleases found throughout bacteria. Independent of the trajectory towards adaptive immunity, TnpB nucleases have also recurrently given rise to TnpB-like nuclease-dead repressors (TldRs), a family of programmable RNA-guided transcription factors, though the physiological roles of most TldR clades remain unclear. Recently, we identified a TldR clade associated with bacterial ABC transporter operons, but this clade has not been characterized in any native host, leaving the biological significance of its predicted regulatory function unknown. Here, we demonstrate direct *in vivo* repression of the oligopeptide permease (*opp*) binding protein OppA in *Enterococcus faecalis*, implicating TldR in shaping the substrate repertoire of bacterial peptide import machinery. Phylogenetic and comparative genomic analyses reveal that bacterial genomes typically encode multiple, structurally conserved but functionally diversified OppA paralogs, suggesting that TldR-mediated repression could enable selective tuning of transporter composition. To define the structural basis of this regulatory specificity, we used cryo–electron microscopy to capture an *oppF*-associated TldR in multiple functional states, revealing a conserved bilobed architecture and a TAM recognition mechanism inherited from TnpB ancestors. Together, these findings define the structural, genetic, and evolutionary basis of a widespread RNA-guided regulatory system domesticated from mobile genetic elements to tune peptide transport in bacteria.

## INTRODUCTION

Transposons have served as direct evolutionary reservoirs for CRISPR-Cas immune effectors^1^, with TnpB and IscB established as the ancestral predecessors of Cas12 and Cas9, respectively^2–5^. TnpB is among the most abundant gene families in nature, encoded within IS*200*/IS*605* and IS*607* transposons as an RNA-guided endonuclease that promotes transposon maintenance and retention^6–8^. TnpB functions with an adjacently encoded ωRNA comprising a conserved scaffold and a ~16-nt guide derived from the transposon-flanking sequence, which directs TnpB-ωRNA to recognize and cleave the vacant donor joints formed upon transposon excision^9,10^. Repeated exaptation events of TnpB, coupled with ωRNA diversification into distinct gRNAs with varied requirements and target-adjacent motif (TAM) specificities, gave rise to Cas12 nucleases^4,5,9^, demonstrating that TnpB scaffolds can be repurposed for functions far beyond transposon maintenance.

This evolutionary pattern extends beyond CRISPR-Cas immunity, with the emergence of nuclease-dead variants repurposed for RNA-guided transposition^11,12^, positioning TnpB as a recurring reservoir for novel RNA-guided molecular functions. Using a custom bioinformatics pipeline, we recently reported the discovery of numerous nuclease-inactivated TnpB-like proteins bearing progressive RuvC active site mutations yet retaining RNA-guided DNA binding activity^13^. Bioinformatic analyses predicted that the gRNA-matching DNA targets overlap promoter regions of host genes, suggesting a role in transcriptional regulation. Heterologous experiments confirmed this, demonstrating that these proteins sterically occlude RNA polymerase to repress transcription — defining a class of naturally evolved RNA-guided transcription factors we named TldRs (TnpB-like nuclease-dead repressors)^13^. We experimentally characterized three major TldR clades: one associated with a prophage-encoded *fliC* gene in *Enterobacteriaceae*, another associated with a host-encoded *csrA* gene in gut-commensal Firmicutes, and a third associated with a host-encoded *oppF* gene in *Enterococcus*^13^. The *fliC*-associated TldR was subsequently shown to natively remodel flagellar composition, altering motility and reducing host immune recognition^14^. Whether *oppF*-associated TldRs natively repress their predicted target, and what the biological consequences of this activity might entail, remained entirely unexplored.

ATP-binding cassette (ABC) transporters are ubiquitous, membrane-bound protein complexes that move substrates in and out of cells and are responsible for nutrient uptake, drug resistance, toxin efflux, and virulence factor transport, among other roles^15,16^. Oligopeptide permease (Opp) transporters, an ABC transporter subtype, typically consist of five components: OppA, a periplasmic substrate-binding protein (SBP) that captures oligopeptides and delivers them to the OppBCDF transmembrane complex, which drives ATP-dependent import into the cytoplasm^15–17^. These transporters intersect with nutrient sensing, intracellular signaling, and virulence in pathogenic organisms, necessitating tight transcriptional regulation^16,17^. Moreover, OppA paralogs are often present in multiple copies within bacterial genomes, implying diverse roles in distinct substrate binding and signaling^15^. Strikingly, *oppF*-associated TldRs are predicted to target the promoter regions of their cognate *oppA* genes^13^, positioning ABC transporter regulation as a candidate physiological role for this TldR clade.

Here, we define the molecular and evolutionary basis of *oppF*-associated TldR function using transcriptomic, comparative genomic, and structural analyses. Working in native *E. faecalis*, we demonstrate that TldR directly represses *oppA* expression in vivo, establishing its role as a functional RNA-guided regulator of ABC transporter genes. Unexpectedly, we also observe upregulation of several paralogous *opp* genes upon TldR overexpression, highlighting a more intricate regulatory network between peptide transporters. Cryo-electron microscopy structures of *oppF-*associated TldR help elucidate a nearly universally conserved, TAM-dependent DNA binding mechanism across TldR clades. These findings further establish TnpB domestication as a rich source of evolutionary diversity, expanding the known repertoire of RNA-guided cellular functions.

## RESULTS

### OppF-associated TldR represses oppA expression in Enterococcus

We previously demonstrated that an *oppF-*associated TldR homolog from *E. faecalis* isolate WE0851 (*Efa_1_*TldR) represses reporter gene expression in an *E. coli* heterologous system using an engineered target site^13^. However, we reasoned that elucidating the native biological consequences of *oppF*-associated TldR activity would require working directly in *Enterococcus*, the predominant natural host of this TldR clade (**Fig. S1a**). Among *Enterococcus* species, *E. faecalis* is one of the most extensively studied — an opportunistic human pathogen that colonizes the gastrointestinal tract and is a leading cause of hospital-acquired infections, including bacteremia, urinary tract infections, and endocarditis, driven by its intrinsic and readily acquired antibiotic resistance^18–20^. This ecological versatility is linked in part to rapid metabolic and transcriptional adaptation across host niches, including the gastrointestinal tract, urinary tract, and bloodstream^20^, raising the possibility that systems such as Opp transporters contribute to this adaptability. The predicted target sites of *oppF*-associated TldRs are consistently located upstream of *oppA* genes (**Fig. S1b**), yet this had never been tested in a native host. We therefore worked in the clinical isolate *E. faecalis* V583, a genetically tractable host with established tools for genetic manipulation^21–23^, which encodes a TldR homolog that is 99.7% identical to *Efa_1_*TldR at the amino acid level, an identical predicted gRNA sequence, and a conserved *oppA* upstream region.

To uncover the regulatory function of TldR, we sought to analyze transcriptome-wide changes under different genomic contexts in *E. faecalis* V583 via RNA-seq (**Fig. 1a**). We started by generating two recombineered strains: a ‘watermarked’ control strain (hereafter WT), in which a neutral 10-bp sequence was inserted downstream of the gRNA, to control for recombineering-induced effects, and a Δ*tldr*-gRNA deletion strain, both generated using a derivative of the scarless recombineering pLT06 vector^22^. Correct genotypes were confirmed by whole genome sequencing (**Fig. S1c**), and RNA-seq revealed all components of the operon to be actively expressed in the WT strain, albeit with TldR at a lower level than the rest of the operon (**Fig. 1b; Table S4**).

**Figure 1.**
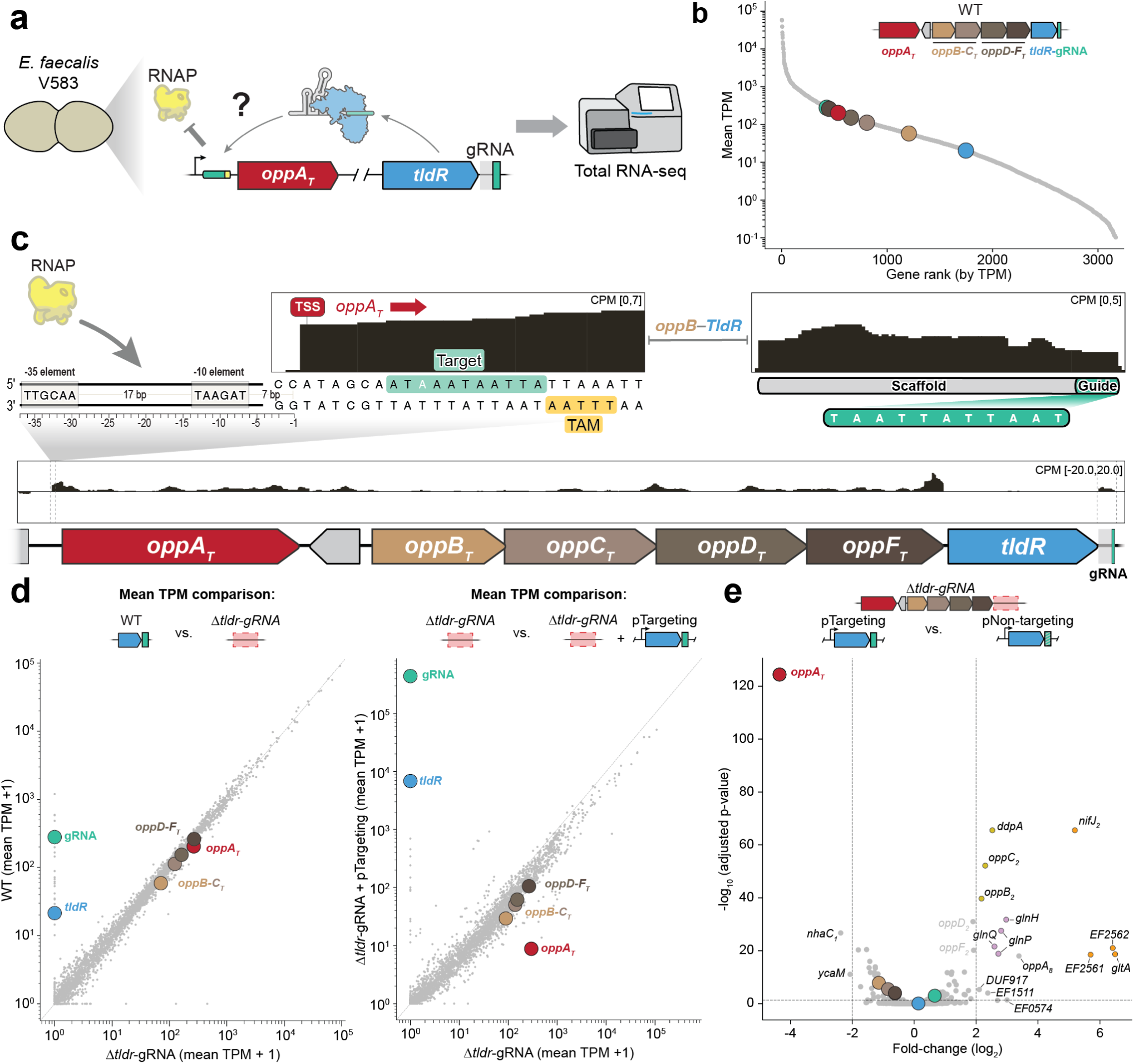
Native TldR function in *E. faecalis*. **a,** Workflow for *E. faecalis* V583 total RNA-seq. Native genetic and transcriptomic experiments were carried out to test native TldR-mediated repression of its predicted target. **b,** Mean transcriptome-wide expression across three biological replicates of the WT control strain. Gene rank (x-axis) versus mean TPM (y-axis, log_10_ scale) for genes in *E. faecalis* V583. Features with mean TPM < 0.1 were excluded from visualization. **c,** RNA-seq coverage tracks over the *oppA-oppBCDF-tldR*-gRNA operon in the WT control strain, normalized to counts per million (CPM), showing sense (top sub-track) and antisense (bottom sub-track) strands. The operon’s continuity is interrupted by a ~100 amino acid protein of unknown function (DUF3899) encoded antisense to the rest of the operon (grey). Top-left: zoom-in of the *oppA* upstream region, highlighting the TSS, BPROM predicted promoter, guide RNA target site (green), and TAM sequence (gold). Top-right: zoom-in of the gRNA coverage, showing the scaffold (grey) and guide sequences (green). This target site position revises the targeting architecture previously proposed for *Efa_1_*TldR, in which the promoter was computationally predicted to overlap the target^13^. RNA-seq mapping here instead places the true TSS upstream of the target, showing that the target site falls within the transcribed portion of *oppA_T_*. **d,** Scatter plot comparison of mean transcriptome-wide expression between the Δ*tldR*-gRNA strain and the WT control strain (left), and Δ*tldR*-gRNA and the pTargeting TldR-gRNA strain (induced, right). Both axes show mean TPM + 1 (log_10_ scale). **e,** Differential expression between targeting and non-targeting gRNA conditions, both ectopically expressed in the Δ*tldR*-gRNA background. Volcano plot showing log_2_ fold-change (x-axis) versus adjusted p-value (y-axis). Dashed lines indicate significance thresholds (|log_2_FC| > 2, padj < 0.05). Highlighted genes of the *oppF-TldR* operon are shown as in (c). Genes highlighted in gold, pink, and orange are operonic.

Inspection of RNA-seq coverage tracks over the *tldR*-associated *oppA* (hereafter *oppA*_T_) locus in the WT strain revealed a clear transcription start site (TSS) positioned 7-bp downstream of a BPROM-predicted –10 element, consistent with a canonical sigma factor binding architecture^24,25^ (**Fig. 1c**). Coverage over the gRNA region revealed clear edges demarcating the boundaries of the scaffold and guide sequence, consistent with our previously reported guide length and seed sequence^13^. Interestingly, the predicted gRNA-matching target site, with a cognate adjacent TAM, is positioned downstream of the *oppA_T_* TSS, which contrasts with the putative targeting architecture observed in *E. faecium*, where the TAM and target site directly overlap a predicted promoter upstream of the *oppA_T_* TSS^13,26^, suggesting natural variation in the specific mechanism of RNAP occlusion and gene repression across the *oppF*-associated TldR clade.

To assess the regulatory effects of TldR on *oppA*_T_ expression, we first examined RNA-seq data from the Δ*tldr*-gRNA strain. Consistent with the modest native expression of TldR under standard culturing conditions, deletion of TldR and its gRNA caused virtually no genome-wide transcriptional changes, including no significant *oppA_T_* de-repression (**Fig. 1d**). We therefore ectopically overexpressed TldR and its gRNA from an agmatine-inducible plasmid^23^ in a Δ*tldr*-gRNA background and repeated RNA-seq analyses. Strikingly, within this genetic background, we observed both robust TldR-gRNA induction upon agmatine treatment, as well as severe (~40-fold) concomitant *oppA_T_* repression (**Fig. 1d**). When we performed RNA-seq with a non-targeting gRNA control and compared transcriptome-wide expression changes, *oppA_T_* was the most significantly repressed gene (**Fig. 1e**). No agmatine-dependent transcriptional effects were observed, and expression of the downstream *opp_T_* genes remained largely unchanged (**Fig. S2a**), consistent with the hypothesis that *oppA_T_* expression is driven by a dedicated promoter distinct from that controlling the expression of the rest of the operon (**Fig. S2b**). Together, these data demonstrate that TldR functions as a direct RNA-guided transcriptional repressor of *oppA_T_* in its native *E. faecalis* host.

Unexpectedly, we also observed differential expression of several other genes upon TldR-gRNA overexpression (**Fig. 1e**). Two other genes were modestly repressed (*nhaC_1_* and *ycaM*), whereas several genes were significantly up-regulated, including a paralogous Opp operon encompassing *ddpA*, *oppB_2_*, *oppC_2_*, *oppD_2_*, and *oppF_2_*, direct paralogs of the *oppA_T_*-*oppF_T_* components (**Fig. 1e**). An orphan *oppA* paralog (*oppA_8_*) was also upregulated, alongside the *glnP-glnH-glnQ* (*EF1117–EF1120*) operon, a putative ABC transporter operon with structural homology to a glutamine transporter, and the *nifJ_2_*-*gltA*-*EF2561*-*EF2562* operon, encoding enzymes involved in flavodoxin-mediated electron transfer and glutamate synthesis. Analysis of the upstream regions of all differentially expressed genes revealed no TAM-adjacent gRNA complementarity other than at *oppA_T_* (**Fig. S2c**), arguing against off-target TldR activity, and the observed transcriptional changes are also unlikely to reflect agmatine-induced artifacts (**Fig. S2a**). These effects thus extend beyond direct *oppA_T_* repression, revealing compensatory expression changes across paralogous peptide transporter loci in *E. faecalis*.

### Comparative genomics suggest TldR-mediated rewiring of ABC transporter regulation

OppA, the periplasmic SBP of the Opp system introduced above, engages the OppBCDF membrane complex only transiently, with productive docking driven by ligand-induced closure and coupled to ATP hydrolysis and substrate transport (**Fig. 2a**)^15,16,27,28^. We hypothesized that TldR-mediated *oppA* repression might exert its biological effects by freeing the OppBCDF complex to engage alternative SBPs, prompting us to investigate the diversity and functional potential of OppA paralogs encoded elsewhere in the V583 genome.

**Figure 2.**
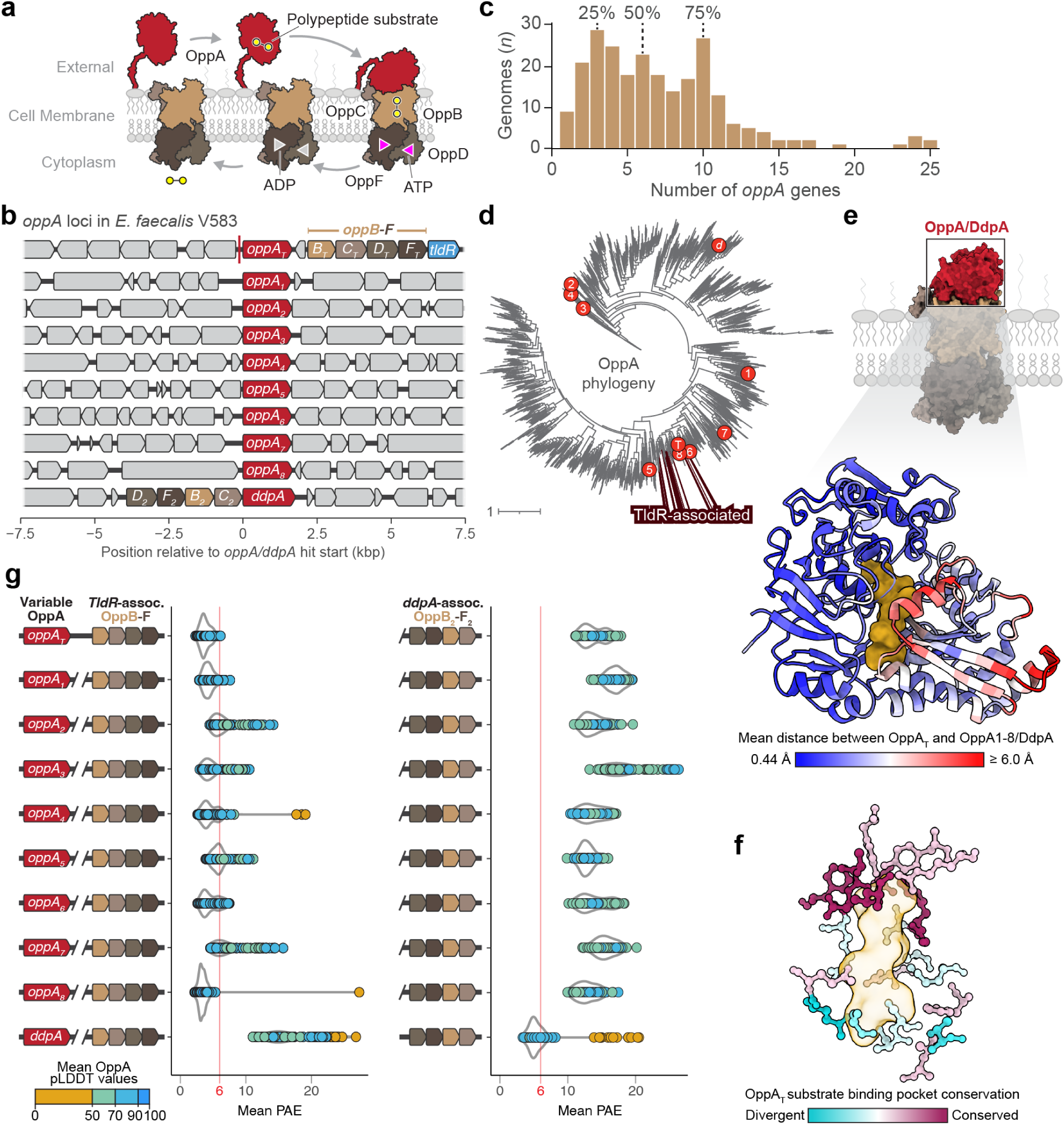
Comparative genomics and structural diversification of OppA paralogs. **a,** Schematic of the oligopeptide permease (Opp) transport cycle. The periplasmic substrate-binding protein OppA captures extracellular oligopeptides and delivers them to the OppBCDF membrane complex, where ATP hydrolysis drives substrate translocation into the cytoplasm. **b,** Genomic organization of *oppA* loci in *E. faecalis* V583. Multiple *oppA* paralogs are distributed across the genome, including the TldR-associated *oppA_T_* and additional paralogs found either adjacent to distinct *oppBCDF* operons or as orphan genes in diverse genomic contexts. **c,** Histogram showing the distribution of *oppA* copy number across complete bacterial genome assemblies with dashed lines indicating the median and interquartile range (25th and 75th percentiles). **d,** Phylogenetic tree of OppA homologs, with TldR-associated homologs forming a distinct clade (highlighted), consistent with functional specialization. **e,** Structural divergence among OppA paralogs. AlphaFold 3-predicted structures of V583 OppA paralogs were superimposed, and pairwise distances between corresponding Cα atoms were calculated relative to OppA_T_ and mapped onto the OppA_T_ structure. **f,** Variation in predicted substrate-binding pockets across OppA paralogs visualized as sequence conservation mapped onto the OppA_T_ predicted structure, consistent with diversification of ligand specificity. **g,** Predicted interaction specificity between OppA paralogs and each of the two OppBCDF transporter assemblies encoded in the V583 genome (left, *tldR*-associated OppB–F_T_; right, *ddpA*-associated OppB–F_2_). Mean predicted aligned error (PAE) values derived from AlphaFold 3 models of OppA–OppBCDF complexes are shown as per-residue-pair values (dots), colored by the mean pLDDT of each pair. A mean PAE of 6 (red vertical line) denotes the threshold for high-confidence interactions, where lower values reflect higher confidence, revealing a partitioned interaction landscape in which subsets of OppA paralogs show selective compatibility with distinct transporter assemblies.

We identified a second *oppABCDF* operon and eight additional standalone *oppA* homologs in diverse genetic contexts (**Fig. 2b**), revealing a substantial combinatorial capacity for diversifying Opp-mediated peptide uptake. A more systematic analysis of fully assembled genomes further revealed that bacteria typically encode multiple *oppA* paralogs^15^, with a median of 6 (IQR = 3-10) in our survey of complete genome assemblies (**Fig. 2c**), and that these homologs are quite diverse at the amino acid level (**Fig. 2d; Fig. S2d**). Structural comparisons of OppA paralogs revealed a high degree of structural similarity, apart from the surface-exposed regions (**Fig. 2e**), preserving a conserved overall fold. Strikingly, the predicted substrate-binding pocket exhibited substantial variability across paralogs (**Fig. 2f**), supporting a model in which distinct OppA proteins encode differentiated substrate specificities that can be selectively engaged by the associated OppBCDF membrane transporter.

To explore whether distinct OppA paralogs preferentially engage specific OppBCDF transporters, we used AlphaFold 3 to model pairwise complexes between each OppA paralog and both OppBCDF assemblies encoded in the *E. faecalis* V583 genome. We then quantified model confidence using predicted aligned error (PAE) values across the OppA–transporter interface, reasoning that lower PAEs reflect more favorable and specific interactions. This analysis revealed that the OppA_T_ homolog regulated by TldR exhibited amongst the strongest predicted interactions with its cognate OppBCDF_T_ complex, while several additional paralogs (e.g., OppA_1_, OppA_6_, OppA_8_) also showed comparably strong interaction confidence with this transporter (**Fig. 2g**). In contrast, *ddpA* — which is encoded adjacent to a distinct oppBCDF operon — displayed weak predicted interaction with the TldR-associated transporter but strong interaction confidence with its cognate complex (**Fig. 2g**).

Together, these patterns suggest that most OppA paralogs are broadly compatible with the TldR-associated transporter, whereas *ddpA* is preferentially matched to its own, alternative assembly. This asymmetry is consistent with a largely shared, but partially partitioned, interaction landscape that could be dynamically reshaped by TldR-mediated *oppA_T_* repression. To define how TldR enables this regulatory switch at the molecular level, we next sought to determine the cryo-EM structure of the TldR complex.

### Overall structure of the *Efa_1_*TldR–gRNA–DNA ternary complex

Given the near-identity between V583 and *Efa_1_*TldR previously established, we used *Efa_1_*TldR to determine the structural mechanism of RNA-guided target recognition. We determined the cryo-EM structure of a ternary complex comprising *Efa_1_*TldR bound to its gRNA and a 42-bp double-stranded DNA (dsDNA) substrate containing a 5′-TTTAA-3′ TAM sequence and 9-bp target region, consistent with prior functional analyses showing that 9-bp base-pairing is sufficient for RNA-guided repression^13^ (**Fig. 3a; Fig. S3; Table S5**). Initial cryo-EM datasets collected on holey carbon grids exhibited a strong preferred particle orientation (**Fig. S4**), which we overcame by collecting data on graphene oxide–coated grids (**Methods**), enabling a structure at 3.1 Å resolution (**Fig. 3b; Fig. S3c–h**). The resulting map allowed modeling of nearly the entire complex, with the exception of the Lid motif and C-terminal domain (CTD) (**Fig. 3c,d; Fig. S5**). DNA density was well resolved at the TAM and RNA–DNA heteroduplex, whereas distal regions of both strands were not visible, indicating substantial flexibility (**Fig. 3a,d–f; Fig. S5**). A minor fraction of particles formed a head-to-head dimer whose physiological relevance remains unclear (**Figs. S3g and S6**).

**Figure 3.**
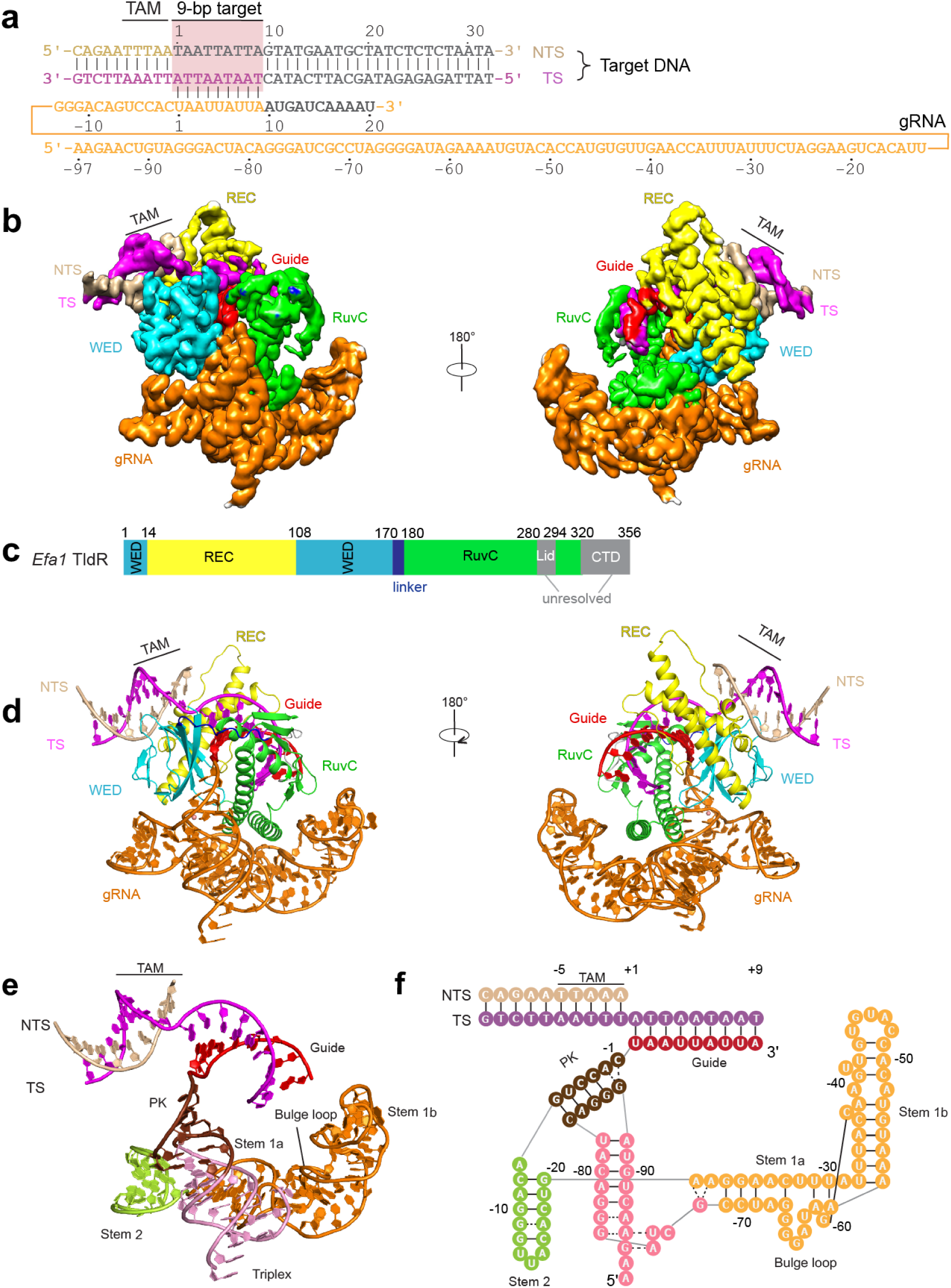
Overall structure of *Efa_1_*TldR–gRNA–DNA ternary complex. **a,** Schematic of the gRNA and target DNA substrate used for cryo-EM analysis, highlighting the TAM and 9-bp target region (see **Methods** for substrate design). Regions not resolved in the final reconstruction and model are shown in grey. TS, target strand; NTS, non-target strand. **b,** Cryo-EM density map of the TldR–gRNA–DNA ternary complex. **c,** Domain organization of *Efa_1_*TldR. The Lid motif and CTD (grey) were not resolved in the cryo-EM structure. **d,** Atomic model of the TldR–gRNA–DNA ternary complex in two views, colored as in **a–c**. **e,** Structure of the gRNA and target DNA, with individual RNA elements annotated. PK, pseudoknot. **f,** Secondary structure representation of the gRNA and target DNA, depicted as in **e**.

Overall, *Efa1*TldR adopts a bilobed architecture comprising an N-terminal recognition lobe (REC and WED domains) connected to a C-terminal RuvC domain (**Fig. 3c,d**). The REC and WED domains closely resemble those of TnpB^29–31^ with modest local differences, including a shortened α2 helix in the REC domain (**Figs. S7a and S8**). In contrast, the RuvC domain exhibits clear degeneration, consistent with prior bioinformatic analyses^13^. The C-terminal region is truncated and largely disordered, and the catalytic DED triad is disrupted, with substitution of the first acidic residue and loss of density for the third (**Fig. 3d; Figs.S5c and S7b**). These distinct features are consistent with evolutionary loss of nuclease activity while preserving the overall TnpB/TldR architectural organization.

The gRNA is bound along a positively charged surface spanning the RuvC and WED domains (**Fig. 3d; Fig. S9**) and comprises a 9-nt guide and a 97-nt scaffold organized into conserved structural elements including a triplex, two stems, and a pseudoknot (PK) (**Fig. 3e,f**). A well-defined globular density at the PK center is consistent with a coordinated Mg^2+^ ion that stabilizes the gRNA core (**Fig. S7c**)^32,33^. A notable feature is the conformation of Stem 1b: whereas this region becomes flexible upon heteroduplex formation in IS*Dra2* TnpB and has been linked to trans-cleavage activity^29^, Stem 1b in TldR remains well ordered despite limited protein contacts and adopts a conformation intermediate between TnpB binary and ternary states (**Fig. 3e,f; Fig. S7a**). This conformation is stabilized by intramolecular RNA interactions that tether Stem 1b to Stem 1a, including stacking and base-pairing contacts within a bulged loop, and likely restrict conformational rearrangements associated with nuclease activation (**Fig. S7d**). Stem 2 is primarily engaged by the WED domain through electrostatic interactions with the RNA backbone, providing additional stabilization of the scaffold (**Fig. 3e,f; Fig. S9**). Together, these features define a compact and conformationally constrained RNA scaffold that supports target engagement, setting the stage for TAM-dependent DNA recognition.

### TAM and heteroduplex recognition

The 5′-TTTAA-3′ TAM duplex is accommodated in a groove formed by the REC and WED domains, defining a DNA recognition module inherited from the TnpB ancestor (**Figs. 3b,d and 4a**). This interaction is anchored by Q116, which inserts into the major groove and forms base-specific contacts across the TAM, with additional contributions from L75 and S74 and a network of conserved surrounding residues that stabilize the duplex (**Fig. 4a–c; Fig. S8**). Consistent with this structural model, mutational analysis using an *in vitro* transcription inhibition assay showed that substitution of either Q116 or L75 strongly impaired TldR-mediated repression, with a charge-introducing mutation at L75 largely abolishing activity (**Fig. 4d; Fig. S7e**). Together, these data identify Q116 and L75 as key determinants of TAM recognition.

**Figure 4.**
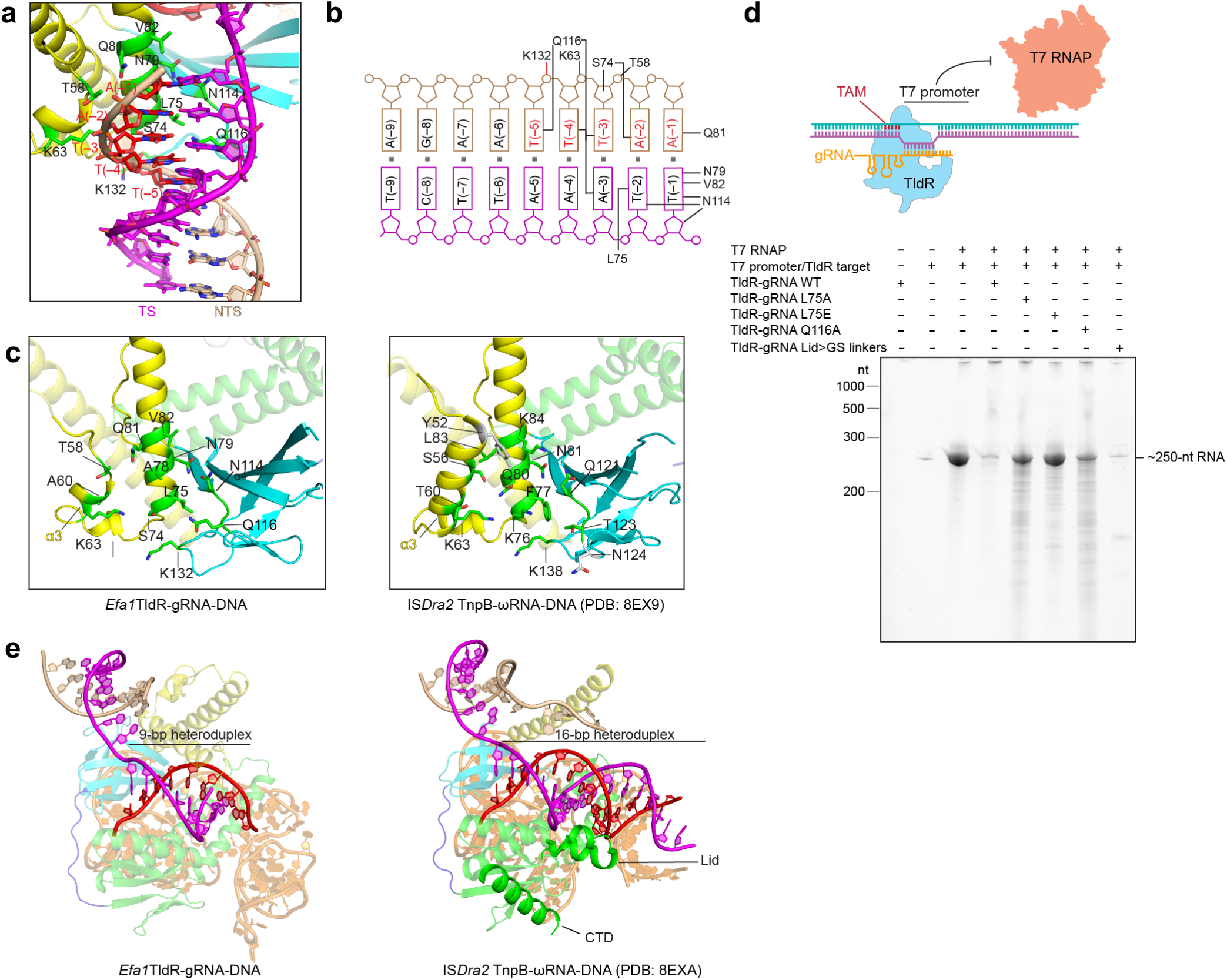
TAM recognition and RNA–DNA heteroduplex architecture in the *Efa1*TldR ternary complex. **a,** Close-up view of TAM recognition by TldR, highlighting interactions between the DNA duplex and the REC and WED domains. **b,** Schematic summary of base-specific and backbone interactions within the TAM duplex. **c,** Comparison between *Efa1*TldR-mediated TAM recognition (left) and IS*Dra2* TnpB-mediated TAM recognition (right). Conserved and divergent features are indicated. **d,** *In vitro* transcription inhibition assay showing TldR-mediated repression of T7 RNAP. The schematic (top) visualizes the TldR–gRNA binding site within the T7 promoter, and the urea-PAGE gel (bottom) shows reduced full-length transcript formation in the presence of the WT TldR–gRNA complex. Results are shown for various TldR perturbations; similar data were obtained from three biological replicates. **e,** Comparison of the RNA–DNA heteroduplex architecture between *Efa_1_*TldR (9-bp heteroduplex) and IS*Dra2* TnpB (16-bp heteroduplex). The Lid motif and CTD, ordered in the TnpB structure, are disordered and unresolved in *Efa_1_*TldR.

Compared with IS*Dra2* TnpB^29,30^, the TldR TAM recognition network is less elaborate. In TldR, the terminal base pair is contacted primarily by L75, V82, N79, Q81, and N114, whereas TnpB engages additional TAM-interacting residues, including Y52 and L83, contributed in part by an elongated α3 helix within the REC domain (**Fig. 4c**). Several key TAM-recognition residues in TnpB are also altered in TldR, including substitutions at positions equivalent to Q80, T60, and K84, consistent with a reduced and reorganized recognition interface (**Fig. 4c**). The RNA–DNA heteroduplex is accommodated within a cleft formed by the REC, WED, and RuvC domains (**Figs. 3b,d and 4e**), with the TAM-proximal base pair stabilized by stacking interactions with aromatic residues Y307 and W311 that are highly conserved across TldR and TnpB homologs (**Figs. S8 and S9**). Additional contacts from L4 and L167, which are unique to *oppF*-associated TldRs, further reinforce this region. The distal region of the 9-bp heteroduplex is stabilized primarily through electrostatic interactions between the nucleic acid backbone and the surrounding protein surface (**Fig. S9**).

The Lid motif — required for substrate loading into the RuvC active site in TnpB and Cas12 enzymes^29,34,35^ — is notably disordered in *Efa* TldR, functionally dispensable for transcriptional repression activity, and poorly conserved across *oppF*-associated TldRs (**Fig. 4d,e; Fig. S8**). The CTD is likewise unresolved in the structure (**Fig. 4e; Fig. S5c**), although its deletion led to protein precipitation during purification, suggesting a structural role in maintaining RuvC integrity.

Together, these structural and mutational data define Q116, L75, and their surrounding network of conserved residues as the core determinants of TAM recognition in TldR, while also revealing a simplified and reorganized TAM-binding interface relative to its TnpB ancestor.

### Structure of the *Efa1*TldR–gRNA binary complex

To gain insights into structural rearrangements induced by target engagement, we determined the cryo-EM structure of the TldR–gRNA binary complex. Reconstructions were obtained for a dimeric assembly at 4.68 Å resolution and for a single protomer at 3.37 Å resolution (**Figs. S10a and S11; Table S5**). Consistent with SEC analysis, the complex primarily adopts a dimeric state in solution (**Fig. S3a**); owing to conformational flexibility of the dimer, the dimeric reconstruction was obtained at lower resolution (**Fig. S11d–j**). Dimerization appears to be mediated by RNA–RNA interactions involving the guide regions, which bring the two TldR molecules together (**Figs. S10a and S11**), although the interface could not be resolved in detail. Most protein domains are not resolved in the binary complex, indicating substantial conformational flexibility. Only the α1 and α2 helices of the RuvC domain are clearly visible, likely stabilized by gRNA binding (**Fig. S10a**). In contrast, the gRNA scaffold closely resembles that observed in the ternary complex and could be readily fitted using the ternary model (**Fig. S10a–d**), suggesting that much of the RNA architecture is pre-organized prior to DNA binding.

A key difference lies in the conformation of the guide region. Unlike TnpB and Cas12-family effectors that typically exhibit a pre-organized seed region^29,36,37^, the guide segment of *Efa1*TldR adopts a stem-loop structure in the binary complex (**Fig. S10a,c**). The stem-loop occupies a position near the WED domain and overlaps with the TAM binding site observed in the ternary complex (**Fig. S10d**). Thus, formation of the DNA-bound ternary complex likely requires remodeling of the guide region to enable TAM recognition and RNA–DNA heteroduplex formation.

## DISCUSSION

TnpB-derived TldR proteins are emerging as significant regulators of host transcription, with consequences that extend well beyond their ancestral role in transposon maintenance (**Fig. 5**). Our findings add to a growing body of evidence — from phage-mediated remodeling of flagellar composition^13,14^ to the RNA-guided transcriptional tuning of ABC oligopeptide transporter expression shown here — that exapted TnpB-like proteins have been repeatedly co-opted to reshape the transcriptomic landscape of their hosts. The widespread occurrence of multiple, functionally diversified OppA paralogs across bacterial genomes further suggests that TldR and OppA diversification are evolutionarily linked, establishing a broader regulatory network controlling peptide transporter composition (**Fig. 5**). Our transcriptomic data show that repression of a single OppA variant shifts the relative contribution of remaining paralogs, and since distinct OppA paralogs likely differ in substrate specificity, TldR-mediated tuning of their expression could directly influence which peptides are preferentially imported to the cell. The broader differential expression we observed in fermentative and anaerobic metabolic pathways further suggests that this regulation extends to cell-wide metabolic shifts, with potential consequences for nutrient sensing and virulence in *E. faecalis* — consequences that will require deeper future investigation.

**Figure 5.**
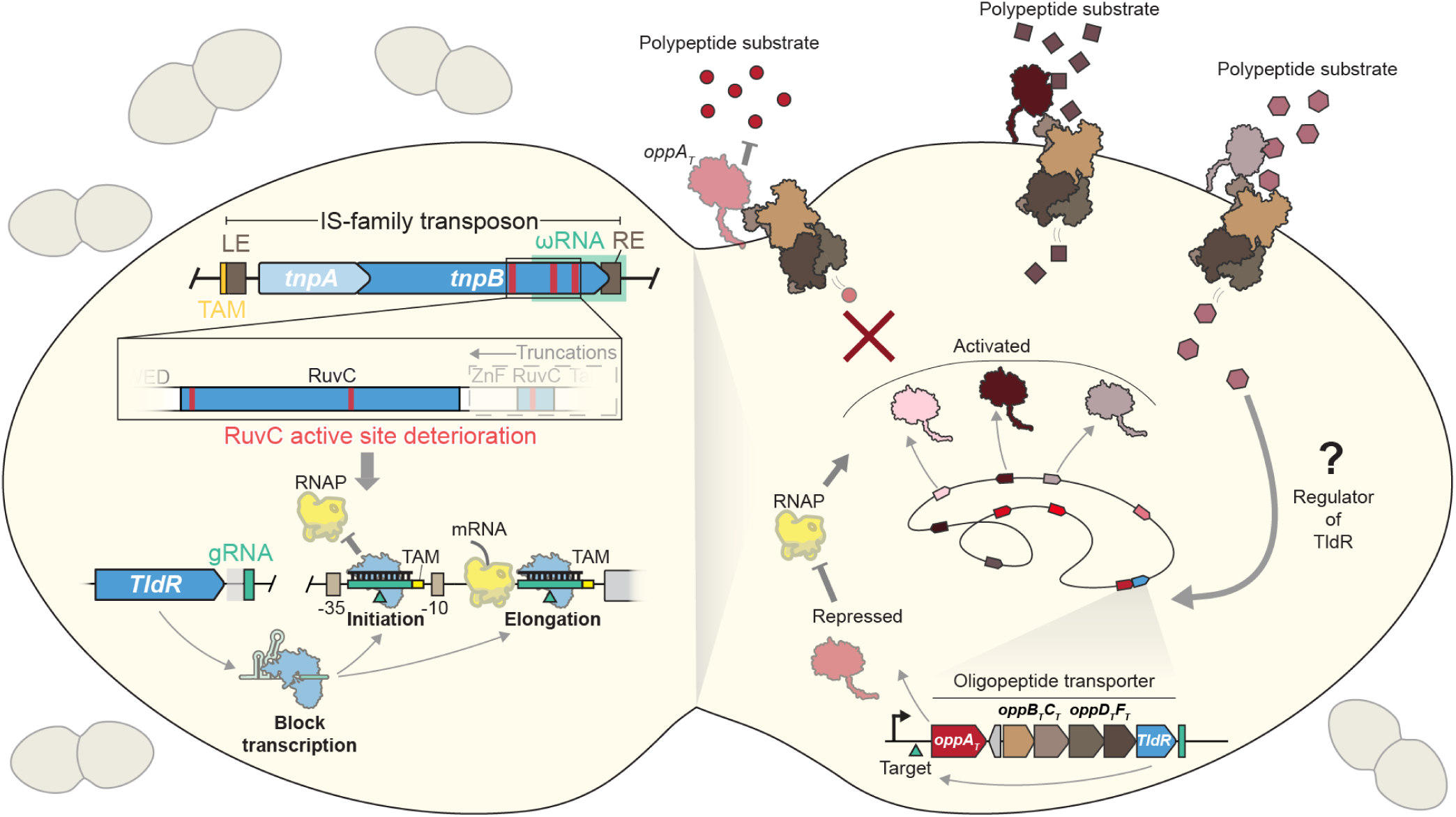
Model for RNA-guided control of ABC transporter composition by TldR. TldR proteins arose repeatedly from the evolutionary repurposing of ancestral, transposon-encoded TnpB nucleases (left), retaining RNA-guided DNA binding while losing DNA cleavage activity through progressive deterioration of the RuvC active site. In their domesticated form, TldRs are guided by a short guide RNA sequence to target sites positioned at or downstream of the promoter, blocking either RNA polymerase binding or its subsequent elongation, and thereby repressing transcription. In *E. faecalis*, this mechanism directly represses expression of a specific *oppA_T_* gene encoding a substrate-binding component of the oligopeptide permease (Opp) transporter. Because bacterial genomes often encode multiple distinct *oppA* homologs, repression of one variant enables alternative substrate-binding proteins to associate with the shared OppBCDF transporter, effectively shifting the spectrum of imported peptides. This regulatory logic likely provides a dynamic mechanism for tuning transporter composition in response to environmental conditions, allowing bacteria to adapt nutrient uptake and associated physiological processes to changing ecological or host-associated contexts. The native signal(s) that induce TldR expression remain unknown.

TldR represents a naturally evolved parallel to engineered CRISPRi^38,39^, likely sharing the core mechanism of RNA-guided steric interference with RNA polymerase, without any associated transcriptional repressor domain. Indeed, just as CRISPRi can be programmed to block either RNAP binding or RNAP elongation depending on target position^40^, the natural variation in target sites we observe across the *oppF*-associated TldR clade — promoter-proximal targeting in *E. faecium*^13,26^ versus downstream, elongation-blocking targeting in *E. faecalis* — suggests that TldR has converged on this same mechanistic flexibility. This indicates that steric exclusion of RNA polymerase by an RNA-guided DNA-binding protein is a particularly accessible and effective regulatory strategy, one achieved independently through TnpB domestication in nature and through deliberate protein engineering by humans. We recently reported a distinct domestication event involving a bacterial Cas12 protein that similarly lost nucleolytic activity and retained RNA-guided DNA binding, but additionally acquired association with a host sigma factor, enabling RNA-guided transcriptional activation^41,42^. In contrast to TldR, that system requires additional regulatory machinery, underscoring that steric RNAP exclusion by TldR represents a minimal and self-contained solution for transcriptional repression. TnpB-to-TldR domestication exemplifies a broader evolutionary pattern in which transposon-encoded proteins are repeatedly co-opted for host regulatory functions, a pattern that encompasses intron splicing, immunoglobulin gene diversification, genome rearrangement, genome defense, and the emergence of CRISPR-Cas adaptive immunity. Collectively, these examples illustrate that transposable elements are not merely genomic parasites but a recurring reservoir of molecular innovation, with TldR representing one of the clearest demonstrations of this principle in the context of programmable gene regulation.

Our structural analyses reveal that the TnpB-to-TldR transition involves not only loss of nuclease activity, but a suite of architectural adaptations that collectively enforce this loss. The Lid motif, required for R-loop sensing and substrate loading, is disordered in TldR, and Stem 1b of the gRNA, whose rearrangement supports trans-cleavage in TnpB, is structurally constrained. One possible explanation for these additional adaptations is that simple catalytic mutations or RuvC truncations may be insufficient to fully silence nuclease activity under all conditions — indeed, DDE-family nucleases can retain catalytic activity when a single catalytic residue is substituted with cysteine in the presence of Mn^2+^ rather than Mg^2+^ ^43^. Notably, *Efa_1_*TldR contains a D185C substitution within the catalytic triad (**Fig. S7b**), suggesting that these additional structural modifications serve as safeguards against unintended nuclease reactivation following domestication.

*Efa_1_*TldR targets a region downstream of *oppA_T_*’s TSS, indicating that it represses transcription by acting as a roadblock to elongating RNA polymerase rather than by occluding the promoter and blocking transcription initiation. Bacterial CRISPRi systems achieve both modes of repression depending on target position — promoter-proximal targeting blocks RNAP binding, whereas targeting within the transcribed region blocks RNAP elongation — with the efficacy of either mode governed by the thermodynamic DNA binding affinity of the effector, particularly the residues that contact the TAM and heteroduplex^40,44,45^. TldR likely operates through a similar roadblock mechanism, retaining the core RNA-guided DNA recognition properties of its TnpB ancestor. Indeed, substitution of either Q116 or L75 in the TAM recognition module is sufficient to strongly reverse transcriptional inhibition. However, *Efa_1_*TldR also possesses a simplified TAM recognition module and a significantly shorter heteroduplex with fewer overall interactions. These features may represent evolutionary adaptations associated with its transition from a nuclease to a host transcriptional regulator. In this context, excessively strong or long-lived DNA binding could be disadvantageous, as it may lead to irreversible repression or reduced regulatory flexibility. A weaker yet sequence-specific interaction may instead enable more tunable and dynamic control of gene expression.

The compact architecture of *oppF*-associated TldR — a small protein, simple gRNA scaffold, and an unusually short 9-nt guide — constitutes a minimalist RNA-guided repression system with potential appeal for synthetic biology applications where small payload size and programmability are priorities. The discovery and functional characterization of *fliC*-, *csrA-,* and *oppF*-associated TldR clades makes clear that additional domestication events almost certainly remain uncharacterized, and that the diversity of RNA-guided regulatory logic encoded in bacterial genomes is broader than currently appreciated. Systematic mining of nuclease-dead TnpB homologs across bacterial genomes is likely to reveal further examples of domesticated RNA-guided regulators with distinct biological roles — and perhaps new tools for programmable gene control.

## METHODS

### Bacterial strains and growth conditions

All strains used in this study are described in **Supplementary Table 1**. In brief, *Enterococcus faecalis* V583 WT and recombineered strains were grown in Todd-Hewitt Broth (THB), Brain Heart Infusion (BHI) broth, or M17 broth as stationary cultures at 30 or 37 °C, depending on the experiment. Cultures of strains carrying either pLT06-derived recombineering vectors or TldR-gRNA overexpression vectors for RNA-seq were also supplemented with 25 μg/mL chloramphenicol.

### Recombineering and genetic manipulations in *E. faecalis* V583

Genetic transformation and recombineering in *E. faecalis* V583 were performed as previously described^21,22^. All plasmids and primers used in this study are described in **Supplementary Tables 2 and 3**. In summary, we cloned derivatives of the temperature-sensitive, markerless genome modification vector, pLT06 (a kind gift from Lynn E. Hancock), with two adjacent ~1-kb fragments homologous to regions flanking the loci of interest (*tldR*-gRNA) via standard molecular biology techniques in *E. coli*, to generate the Δ*tldr*-gRNA recombineering vector. Homology arms were subcloned from the *E. faecalis* T11 genome, which harbors 100% nucleotide identity at the selected loci to *E. faecalis* V583. The control locus pLT06-derivative was cloned similarly, with a 10-bp sequence interrupting the homology fragments, which flank an innocuous region downstream of the predicted gRNA terminator.

Electrocompetent cells and transformations were prepared as previously described^21^. *E. faecalis* V583 was made electrocompetent by growing a stationary overnight culture in 100 mL of M17 supplemented with 0.5 M sucrose and 4% glycine at 37 °C for 20–24 h. The culture was chilled on ice for 10–15 min, then spun at 4,000 x *g* for 10 min at 4 °C. All subsequent steps were conducted in a 4 °C cold room with samples kept on ice. Pellets were washed twice with 1 volume of cold electroporation buffer (0.5 M sucrose, 10% glycerol in Milli-Q water), pelleted as above, and resuspended in 1:100 of the original culture volume in electroporation buffer. Resuspended cells were aliquoted in PCR tubes at 50 μL total volume per tube and flash frozen in liquid nitrogen, then kept at –80 °C.

Electrocompetent cells were mixed with 300–400 ng of Δ*tldr*-gRNA and WT control pLT06-derived plasmid DNA on ice and electroporated by pulsing at 2.5 kV, 25 µF capacitance, and 200 Ω resistance. Samples were immediately recovered in 1 mL of ice-cold M17 supplemented with 0.5 M sucrose, 10 mM MgCl_2_, and 10 mM CaCl_2_. Recoveries were incubated at 30 °C without aeration for at least 2 h. After recovery, electrotransformed cells were plated on BHI-agar plates supplemented with chloramphenicol (25 μg/mL) and X-Gal (80 μg/mL), and incubated at 30 °C for 24–48 h.

Recombineering mediated by Δ*tldr*-gRNA and WT control pLT06-derived vectors was performed as previously described^22^. In brief, transformed blue colonies were screened by colony PCR for the presence of the respective vectors. Screened colonies were grown overnight as stationary cultures at 30 °C in THB Chloramphenicol (25 μg/mL) and diluted 1:100 in fresh THB Chloramphenicol (25 μg/mL). Sub-cultures were grown standing at 30 °C for 2.5 h, shifted to 42 °C for an additional 4–5 h (or until turbid) to induce single-site plasmid integration, then plated on prewarmed THB Chloramphenicol (25 μg/mL) X-Gal (80 μg/mL) agar plates and incubated at 42 °C for 24–48 h. Large blue colonies were screened by colony PCR for integration junctions at the desired locus and grown overnight in THB without antibiotic selection at 30 °C to induce plasmid excision. Overnight cultures were diluted 1:1000 in fresh THB without antibiotic selection and grown at 30 °C overnight, then diluted and plated on MM9YEG agar plates (M9-agar plates supplemented with yeast extract and 1% glucose, 10 mM *p*-Chl-Phe, 120 μg/mL X-Gal) and incubated at 37 °C for 24 h to cure the excised plasmid. Large white colonies were screened by colony PCR, and selected clones were grown in THB without antibiotic at 37 °C overnight in preparation for whole genome sequencing (Plasmidsaurus).

Due to low coverage across independent sequencing runs of the Δ*tldr*-gRNA strain, reads from all runs were subjected to de novo genome assembly, following Plasmidsaurus’s technical documentation. Genome size was estimated using Lrge (v0.2.1)^46^, and multiple subsampled read sets were generated using Autocycler^47^. Independent assemblies were performed with Flye (v2.9.6+)^48^, Hifiasm^49–51^, and Plassembler (v1.8.0+)^52^, and resulting assemblies were reconciled, trimmed, and reoriented using Autocycler and dnaapler^53^. Among the candidate assemblies produced, the top-ranked assembly was excluded due to an artifactual fusion of a native plasmid sequence into the chromosomal contig, likely reflecting erroneous selection of the sole circularized candidate. The final assembly was manually selected based on concordance with the expected *E. faecalis* V583 chromosome size and correct resolution of native plasmids as independent circular contigs. This analysis revealed the absence of the cryptic native plasmid pTEF3, which was further confirmed by the lack of RNA-seq coverage over an artificial pTEF3 reference (**Fig. S1c**).

Vectors for overexpression of TldR-gRNA (with targeting and non-targeting guides) were constructed using an agmatine-inducible expression system analogous to the previously described pAGEnt vector^23^. In brief, an AguR-AguB promoter fragment was subcloned directly from the *E. faecalis* V583 genome into the pAT28 shuttle vector^54^, alongside 3×FLAG-TldR-gRNA, using standard molecular biology techniques. Due to frequent acquisition of spectinomycin resistance, we subcloned the Chloramphenicol resistance gene from pLT06^22^ and swapped it in place of the Spectinomycin resistance gene in the pAT28-Pagu-TldR-gRNA vector. The *E. faecalis* V583 Δ*tldr*-gRNA recombineered strain was electroporated with both targeting and non-targeting TldR-gRNA overexpression variants, as described above, and genotyped for the presence of the plasmid by colony PCR.

### RNA extraction and RNA-seq library preparation

In preparation for total RNA-seq of engineered *E. faecalis* V583 strains, glycerol stocks of each strain were streaked on BHI agar plates (with or without antibiotics, as needed). Three isolated colonies from each streaked strain were grown overnight in 3 mL of M17 media supplemented with 0.5% glucose at 37 °C. All cultures were treated with agmatine, as previously described^23^, by diluting 1:100 in 10 mL of fresh M17 0.5% glucose media with and without 10 mM agmatine and grown at 37 °C for 6 h. After induction, cultures were cooled on ice for 10 min and spun at 4,000 x *g* for 5 min at 4 °C. Cell pellets were washed in 1 mL of PBS, spun as before, and washed cell pellets were flash frozen in liquid nitrogen.

For RNA extraction, cell pellets were thawed on ice and resuspended in 500 μL of Zymo DNA/RNA Shield Buffer and transferred to a 2 mL screw-capped tube on ice, after which around 200 μL of 0.1 mm Zirconia/Silica Beads were added to each sample. Tubes were vortexed and loaded onto a BioSpec Mini-Beadbeater-16. All samples were beaten for 2.5 min at 3,450 rpm for three rounds, with 2 min ice incubation in between rounds. After bead beating, samples were spun down for 10 min at 4,000 x *g* at 4 °C, and ~300 μL of the supernatant were carefully transferred to clean, cold 1.7 mL Eppendorf tubes. RNA was extracted from 100 μL of each lysate with the Zymo RNA Clean & Concentrator kit, eluted in 20 μL of DNase/RNase-free water and stored at –80 °C.

Total RNA-seq libraries were prepared by diluting 100 ng of extracted RNA samples to 15 μL in NEBuffer 2 and heated to 92 °C for 2 min to shear RNA. DNA contamination and 5′-pyrophosphate groups were removed in a single reaction combining RppH (NEB) and TURBO DNase (Thermo Fisher), with SUPERase-In RNase Inhibitor (Thermo Fisher) included to protect the RNA, at 37 °C for 30 min. RNA ends were then repaired with T4 PNK (NEB) in 1x T4 DNA ligase buffer (NEB) for 30 min at 37 °C. Samples were purified over RNA Clean & Concentrator columns (Zymo), and the RNA concentration was measured with a DeNovix spectrophotometer. cDNA libraries were generated from 100 ng of input RNA per sample using the NEBNext Small RNA Library Prep kit with Illumina adapters. Next-generation sequencing was carried out on an AVITI Sequencing System (Element Biosciences) using a 2 × 75 Cloud-break Freestyle kit.

### RNA-seq data analysis and differential expression

RNA-seq reads were processed using cutadapt (v.5.0)^55^ to remove Illumina TruSeq adapter sequences, trim low-quality ends, and exclude reads shorter than 15 nt. Trimmed and filtered reads were aligned to WGS assemblies of strain-specific reference genomes, which were all annotated as the NCBI genome assembly ASM778v1 at a 95% annotation identity using Geneious Prime and indexed to have the same starting coordinates, using bwa-mem2 (v.2.2.1)^56^ in paired-end mode with a minimum seed length of 25 bp (-k 25). SAMtools (v.1.17)^57^ was used to filter for uniquely mapping reads using a MAPQ score threshold of 1, and to sort and index the alignments. IGV was used to visualize and inspect coverage tracks generated using bamCoverage (v3.5.6)^58^. Gene-level read counts were quantified using featureCounts (v.2.0.2)^59^ in paired-end mode with strand-specific counting (-s 1), counting feature types CDS, ncRNA, tRNA, and rRNA. Reads overlapping multiple features were assigned using fractional counting (--fraction). Duplicate gene names were disambiguated by appending chromosome-specific suffixes based on genomic position. Differential expression analysis was performed with pydeseq2 (v.0.5.4)^60^ to calculate fold-change and adjusted p-values (FDR) using the Benjamini-Hochberg procedure for each comparison.

### Phylogenetic analysis of OppA proteins, and quantification of occurrences per genome

To assemble a dataset of OppA homologs, a local copy of the NCBI NR database (retrieved on September 5, 2024) was initially queried with a *tldR*-associated OppA sequence (WP_029692685.1), via Phmmer (E-value < 10^-100^)^61^. An additional search of the NCBI NR database was performed using the same query, via Jackhmmer for three rounds (E-value < 10^-50^). The resulting hits from both searches were concatenated and then clustered at 80% amino acid identity using CD-HIT^62^, before cluster representatives were aligned with MAFFT (EINSI option; four iterations)^63^. The resulting alignment of 1,640 OppA sequences was sparingly trimmed with Trimal (-gt 0.01)^64^, and a phylogenetic tree was constructed from the trimmed alignment using IQ-TREE 2 (-m Q.pfam+R10 -T AUTO -alrt 1000 -abayes -B 1000 --bnni)^65^ and visualized with ITOL^66^.

To assess the number of OppA homologs encoded per genome (**Fig. 2c**), genomes encoding the OppA homologs represented in the **Fig. 2d** tree were retrieved. Only complete genome assemblies were analyzed (n = 264) to ensure OppA quantifications were not under-estimated due to the inclusion of incomplete genomic assemblies. Open reading frames (ORFs) were then predicted using Prodigal^67^, and a profile hidden Markov model representing OppA — an output from the Jackhmmer search described above — was used to search the translated ORFs from each genome.

BPROM^68^ was used to predict promoter regions for *oppA* by scanning the upstream region of the defined *oppA_T_* TSS (**Fig. 1c**). Promotech^69^ was used to predict promoter regions for *oppB* (**Fig. S2b**), by parsing 40-bp windows in the *oppA_T_* – *oppB_T_* intergenic region. Promoter probability was then predicted for each window using a probability threshold of 0.5.

### Bioinformatic predictions of *oppA*-associated TldR targets

Cas-OFFinder was used to predict TldR targets in 2,125 genomic assemblies encoding *oppA-tldR*^70^. Two separate strings encoding the TldR TAM sequence and target sequence were used to query these searches (5’–TTTAANNNNNNNNN–3’ and 5’– NNNNNTAATTATTT–3’), and hits were filtered to exclude any matches with mutations between the guide and target in a TAM-proximal 6-nt seed region. Protein coding genes within 150-bp downstream of these putative TldR targets were then clustered using CD-HIT^62^ — first at 100% amino acid identity to remove duplicates, then at 60% amino acid identity to group orthologs. Cluster representatives were manually annotated, and clusters with overlapping annotations were merged. The resulting sizes and annotations of the largest clusters were then graphed in **Figure S1b**.

To identify TAM + guide sequence matches in promoter regions of differentially expressed genes (**Fig. 1e**), the 150-bp of genomic sequence immediately upstream of the start codon was extracted for each gene meeting significance thresholds (padj < 0.05, |log_2_FC| > 2) using strand-aware coordinates from the feature annotation. Each upstream region was searched on both strands, the annotated sequence and its reverse complement, for a 16-bp composite motif consisting of the 5-bp TAM (TTTAA) immediately followed by the 11-nt guide sequence, with the targeting and non-targeting guide sequences searched separately. A candidate match required a perfect match to the TAM; per-nucleotide identity across the full 16-bp motif was then scored for all candidate matches, and the match with the highest number of matching positions was retained per gene. Genes were ranked by log_2_FC, and per-nucleotide match results (match, mismatch, or no TAM-anchored match) were rendered as a heatmap for each motif.

### Structural prediction analyses of OppA-F proteins

Structures of OppA proteins from *E. faecalis* V583 (highlighted in **Fig. 2b**) were predicted using the AlphaFold 3 webserver^71^. To assess the similarity of these proteins, predicted structures were superimposed, and the distances between corresponding atoms were calculated in ChimeraX using the *contacts* command (radius = 4)^72^, relative to the OppA_T_ prediction. Mean distances of corresponding Cα atoms were then calculated and visualized in **Figure 2e**. Substrates were visualized by superimposing OppA structures from the PDB onto the predicted structure of OppA_T_ (PDB: 5KZT and 3DRF).

To assess the conservation of the OppA substrate binding pocket, a multiple alignment of OppA sequences from *E. faecalis* V583 was constructed using MAFFT (LINSI option). The resulting alignment was used to color the conservation of OppA_T_ residues predicted to contact an OppA substrate (superimposed from PDB: 5KZT).

AlphaFold 3 was also used to predict complexes of *E. faecalis* V583 OppA sequences with the two OppB-F cassettes encoded by *E. faecalis* V583. The confidence scores of these predicted multimers were then assessed to evaluate the likely compatibility of each OppA homolog with the two OppB-F transmembrane units (**Fig. 2g**). Confidence scores were calculated as follows. Inter-protomer heavy-atom contacts were identified as all atom pairs occurring within 4.5 Å between the OppA chain and the OppB-F chains, then collapsed to unique residue pairs. For each contacting residue pair, PAE values were extracted from the JSON file output by AlphaFold 3 in both directions of the asymmetric PAE matrix — once with the OppA residue as the aligned reference, and once with the OppB-F residue as the aligned reference — and averaged to produce a per-residue-pair mean PAE. A global mean PAE was then computed across all contacting residue pairs for each predicted complex. pLDDT values for contacting residues were extracted from the B-factor column of the corresponding PDB file and averaged per residue. Per-residue-pair mean PAE values and pLDDT values were then plotted for each predicted complex.

### Plasmid construction, protein expression and purification

The *Efa_1_tldR* gene was subcloned into the bacterial expression vector pET28-MKH8SUMO (Addgene #79526). The gRNA with T7 promoter and T7 terminator was subcloned into a pUC19 vector for co-expression. The *Efa1*TldR and gRNA were co-expressed in Rosetta(DE3)pLysS cells (Novagen #70956). Notably, although pET28-MKH8SUMO (ColE1-type origin) and pUC19 (modified ColE1/pMB1 origin) belong to the same incompatibility group, both plasmids were maintained under short-term dual antibiotic selection^73^, allowing successful co-expression. Cells were cultured in Terrific Broth (TB) to an OD_600_ of 0.6. Protein expression was induced with 0.3 mM IPTG, and cultures were incubated overnight at 16 °C. Cells were harvested and resuspended in lysis buffer (25 mM Tris-HCl pH 7.6, 500 mM NaCl, 5% glycerol) supplemented with 1 mM PMSF and 5 mM β-mercaptoethanol, then lysed by sonication. The lysate was clarified by centrifugation, and the supernatant was applied to Ni-NTA resin. After extensive washing with lysis buffer containing 30 mM imidazole, target proteins were eluted with lysis buffer containing 250 mM imidazole.

The His-SUMO tag was removed by overnight digestion with SUMO protease at 4 °C. The *Efa1*TldR**–**gRNA eluate was diluted in buffer containing 25 mM Tris-HCl pH 7.6, 75 mM NaCl, then loaded onto a Heparin column (Cytiva) and eluted using a linear NaCl gradient (0.1 to 1 M). Following concentration, proteins were further purified by size exclusion chromatography (SEC) using a Superdex 200 Increase 10/300 GL column (Cytiva) in equilibration buffer containing 25 mM Tris-HCl pH 7.6, 250 mM NaCl, 5% glycerol, and 2 mM DTT. Purified fractions were concentrated and stored at –80 °C.

### Electron microscopy

The 42-bp dsDNA substrate used for cryo-EM spans positions –10 through +8 relative to the first base of the target region, matching the native *Efa_1_* genomic sequence (identical to V583) over this span with one exception: position +9 was engineered from A to T to restore full complementarity with the gRNA. Positions +10 through +32 of the substrate are non-specific filler sequence, included only to complete the DNA duplex for cryo-EM sample preparation, and do not correspond to either the *Efa_1_* or V583 genomic sequence. Full substrate sequences are provided in **Supplementary Table 2**.

Aliquots (3 μL) of the TldR–gRNA binary complex (1.5 mg/mL) and the TldR–gRNA–target DNA ternary complex (1 mg/mL) were applied to glow-discharged (45 s) Quantifoil holey carbon grids (R1.2/1.3, 300 mesh). The grids were blotted for 2.5 s and plunged into liquid ethane using a Thermo Fisher Scientific Vitrobot Mark IV. The TldR–gRNA–target DNA ternary complex (0.05 mg/mL) was also applied to graphene oxide (GO)-coated Quantifoil holey carbon grids (R1.2/1.3, 300 mesh). The GO grids were prepared as described previously^74^, then glow-discharged for 5 s and blotted for 5 s before being plunged into liquid ethane.

Cryo-EM data were collected with a Titan Krios G4 microscope (FEI) operated at 300 kV, and images were collected using EPU at a nominal magnification of 105,000× (resulting in a calibrated physical pixel size of 0.822 Å/pixel) with a defocus range of 0.8–2.0 μm. Three images per hole were collected using EPU with the beam-image shift method. Images were recorded on a K3 electron direct detector in super-resolution mode at the end of a GIF-Quantum energy filter operated with a slit width of 20 eV. A dose rate of 20 electrons per pixel per second and an exposure time of 2.01 seconds were used, generating 40 movie frames with a total dose of ~59.2 e−/Å^2^.

For the TldR–gRNA binary complex, we collected 1,562 micrographs. After screening based on CTF-estimated resolution and manual inspection, 1,531 micrographs were selected for further analysis. For the TldR–gRNA–DNA ternary complex on the carbon grid, 1,859 micrographs were selected from a total of 1,920 collected images. For the TldR–gRNA–DNA ternary complex on the GO grid, 4,232 micrographs were selected from a total of 4,390 collected images.

### Image processing

Movie frames were imported into cryoSPARC Live^75^ for on-the-fly processing during data collection. Motion correction was performed using Patch Motion Correction with a binning factor of 2. Contrast transfer function (CTF) parameters were estimated using Patch CTF. A few thousand particles were initially auto-picked without a template to generate 2D class averages for subsequent template-based particle picking. The auto-picked and extracted particles were screened by 2D classification to exclude false and bad particles that fall into 2D averages with poor features. A subset of the particles (100,000 particles) was used to generate initial models in cryoSPARC. All screened particles were then subjected to heterogeneous refinement using the generated initial models, and particles in each 3D class were further refined by homogeneous refinement.

Selected particles were used to train a Topaz model for automated particle picking^76^ across all micrographs. The picked particles underwent sequential processing in cryoSPARC: 2D classification, *ab initio* reconstruction (2–4 classes), heterogeneous refinement, and Non-uniform Refinement. Results were manually inspected after each step to determine whether to rerun with modifications (e.g., adjusting class numbers) or proceed to the next step. In the final Non-uniform Refinement, we applied per-particle CTF correction. Local Refinement was performed when needed, such as for the protomer of the TldR binary complex. Resolution was estimated using the FSC = 0.143 criterion. Maps were post-processed using DeepEMhancer^77^ to reduce noise levels and used for model building and refinement.

### Model building and refinement

AlphaFold 3 models^71^ were used as initial models. The individual structures were manually fitted onto respective maps as rigid bodies in UCSF Chimera^78^ and manually adjusted in COOT^79^. The model was then fitted in COOT with all-molecule self-restraints to maintain hydrogen bonds and stacking interactions between bases. Finally, refinement of the structure models against corresponding maps was performed using the phenix.real_space_refine tool in Phenix^80^.

### Protein sequence analysis

Sequence alignments were performed using Clustal Omega^81,82^, and alignment diagrams were plotted using ESPript^83^.

### Structural analysis and visualization

Figures were generated using PyMOL, UCSF Chimera^78^ and ChimeraX^84^.

### *In vitro* transcription-blocking assay

The gRNA was designed to target the bottom strand of the T7 promoter with a 16-nt spacer. The target DNA carries a T7 promoter with a TAM inserted upstream. The target DNA templates for *in vitro* transcription were PCR-amplified from synthesized gBlocks (IDT) (sequence details in **Supplementary Table 2**) and cleaned by QIAquick PCR Purification Kit (Qiagen). The template DNA (125 ng) was incubated with or without *Efa1*TldR**–**gRNA variants in the reaction buffer (20 mM Tris pH 7.6, 50 mM NaCl, 5% glycerol, 5 mM MgCl_2_, 2 mM DTT, 2 mM Spermidine) for 30 minutes at 30 °C. Transcription was initiated by T7 polymerase and NTPs according to the manufacturer’s instructions (NEB). Reactions proceeded for 60 minutes at 37 °C and were terminated by adding 20 μL 2X loading buffer (SDS-DNA loading dye, 8 M urea) followed by incubation at 95 °C for 5 minutes. RNA transcripts were resolved by 15% TBE-Urea Gel (Invitrogen) and analyzed by SYBR Gold staining.

### Mutagenesis

Single amino acid mutations were introduced using the QuikChange site-directed mutagenesis method. Multiple amino acid mutations were generated by ligating inverse PCR-amplified backbone with mutation-bearing DNA oligonucleotides using the In-Fusion Cloning Kit (Clontech). All mutations were verified by Sanger sequencing.

### Data availability

Next-generation sequencing data will be deposited in the National Center for Biotechnology Information (NCBI) Sequence Read Archive and Gene Expression Omnibus prior to publication. Datasets generated and analyzed in the current study are available from the corresponding authors on reasonable request. Cryo-EM reconstructions have been deposited in the Electron Microscopy Data Bank under accession nos. EMD-76499 and EMD-76500. Coordinates for atomic models have been deposited in the Protein Data Bank under accession nos. 12JX and 12JY.

### Code availability

Custom scripts used for bioinformatics and RNA-seq data analyses are available upon request.

## ACKNOWLEDGMENTS

We thank T.M. Smith, A.J. Robinson, A.I. Palmieri, and R. Rafat for laboratory support; K.L. Palmer for gifting us both *E. faecalis* V583 and T11 strains; L.E. Hancock for advice on *Enterococcus* handling and methodologies, as well as gifting the pLT06 and pAT28 recombineering and shuttle vectors, respectively; the JP Sulzberger Columbia Genome Center for next-generation sequencing support; and L. F. Landweber for qPCR instrument access. H.C.L. was supported by the National Science Foundation Graduate Research Fellowship Program and the Paul & Daisy Soros Fellowship for New Americans. L.C. was supported by the NIH grants R01GM138675 and R35GM158248, and the NSF Faculty Early Career Development Program (CAREER) Award 2339799. S.H.S. was supported by NSF Faculty Early Career Development Program (CAREER) Award 2239685, a Pew Biomedical Scholarship, an Irma T. Hirschl Career Scientist Award, the Howard Hughes Medical Institute Investigator Program, and a generous startup package from the Columbia University Irving Medical Center Dean’s Office and the Vagelos Precision Medicine Fund.

## AUTHOR CONTRIBUTIONS

A.A.C., R.X., L.C., and S.H.S. conceived the project. A.A.C. performed all *Enterococcus* microbiology experiments and RNA-seq analyses. R.X. prepared cryo-EM samples, designed the transcription blocking assay, and collected and processed cryo-EM data, with input from L.C. H.C.L. assisted with early *Enterococcus* experimental troubleshooting and performed OppA phylogenetic analyses. T.W. performed bioinformatic analyses and assisted with data visualization. D.X. performed transcription blocking assays for protein mutants and contributed to molecular cloning and protein purification. L.C. and S.H.S. supervised the study. A.A.C., R.X., L.C., and S.H.S. analyzed and discussed the data and wrote the manuscript, with input from all authors. The order of authors R.X. and A.A.C. was decided randomly.

## COMPETING INTERESTS

S.H.S. is a co-founder and scientific advisor to Dahlia Biosciences, a scientific advisor to CrisprBits and Prime Medicine, and an equity holder in Dahlia Biosciences and CrisprBits. The remaining authors declare no competing interests.

Correspondence and requests for materials should be addressed to L.C. or S.H.S.

## SUPPLEMENTARY TABLES

**Supplementary Table 1** | Strains used in study.

**Supplementary Table 2** | Oligonucleotides used in this study.

**Supplementary Table 3** | Plasmids used in this study.

**Supplementary Table 4** | Transcriptome-wide transcript abundance (TPM) across RNA-seq conditions.

**Supplementary Table 5** | Cryo-EM data collection, refinement, and validation statistics.

