## Supplementary Figures for "Structural and genetic dissection of RNA-guided gene repression by TnpB-derived transcription factors"

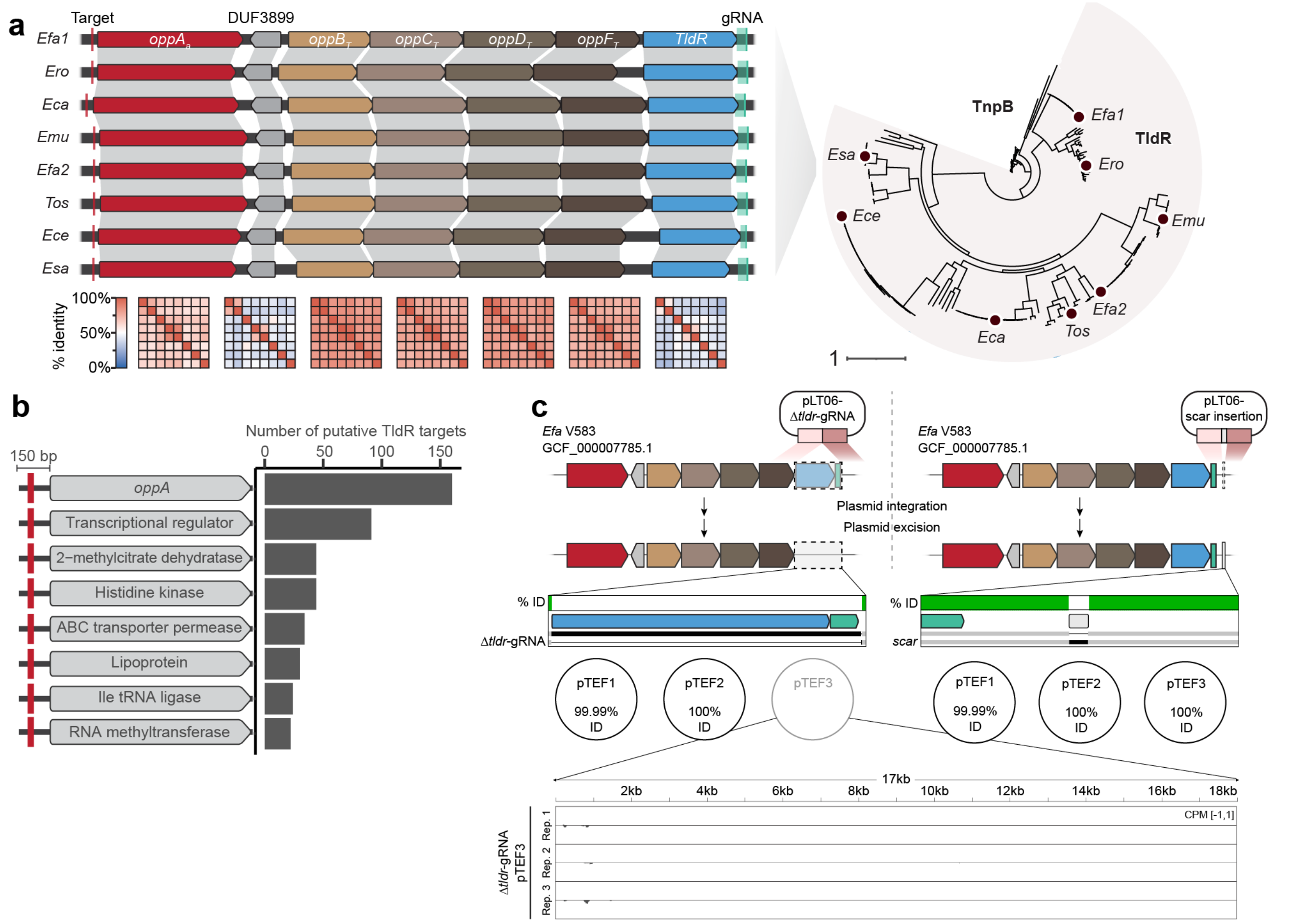


**Figure S1 | *oppF*-associated TldR bioinformatic and *E. faecalis* V583 strain engineering. a,** Left, Genomic organization and phylogenetic context of representative oppF-associated TldR loci. Gene architectures from selected organisms are shown, highlighting the conserved positioning of *tldR* downstream of the *oppABCDF* operon and adjacent to its associated guide RNA (gRNA). The predicted TldR target site is located upstream of the *oppA* open reading frame in each case (red tick), consistent with conserved regulation of the substrate-binding component of the transporter. A small open reading frame (DUF3899) is variably present between *oppA* and *oppB*. Right, phylogenetic tree of TnpB/TldR homologs (adapted from Wiegand *et al.* 2024[13]), with *oppF*-associated TldRs highlighted, indicating their clustering within a distinct clade derived from TnpB ancestors. **b,** Genome-wide prediction of TldR target sites. Putative targets were identified based on guide sequence complementarity and TAM presence, and genes with predicted sites located within 150 bp upstream of their start codons were clustered by functional annotation. The majority of predicted target sites occur upstream of *oppA* genes, which represent the most frequently targeted category. Additional enriched targets (≥20 cluster members) include genes encoding transcriptional regulators, metabolic enzymes, and membrane-associated proteins — the latter including permease subunits of ABC transporter systems unrelated to OppA, which is tallied separately as its own category above — although these occur at substantially lower frequencies. **c,** Top, simplified schematic of the multi-step, scarless recombineering process in *E. faecalis* V583, using pLT06-derived vectors[22] generating a Δ*tldr*-gRNA strain (left) and an innocuous 10-bp insertion downstream of the gRNA as an otherwise isogenic, WT control strain (right). Middle, WGS sequence coverage track insets of the two respective loci previously mentioned. Percent nucleotide identity (% ID) represented as green coverage, indicating 100% identity to the *E. faecalis* V583 NCBI reference genome ASM778v1. White space is indicative of no coverage at that locus (deletion). Native *E. faecalis* V583 plasmids (pTEF1-3) represented as circles, with their respective % ID to the NCBI reference genome. Unexpectedly, we observed loss of the cryptic native plasmid pTEF3 in the Δ*tldr*-gRNA strain, likely attributable to the thermal shock steps inherent to the recombineering workflow[85,86]. Bottom, RNA-seq coverage from a Δ*tldr*-gRNA strain representative sample over an artificially added pTEF3 reference sequence, supporting the loss of the plasmid.

---

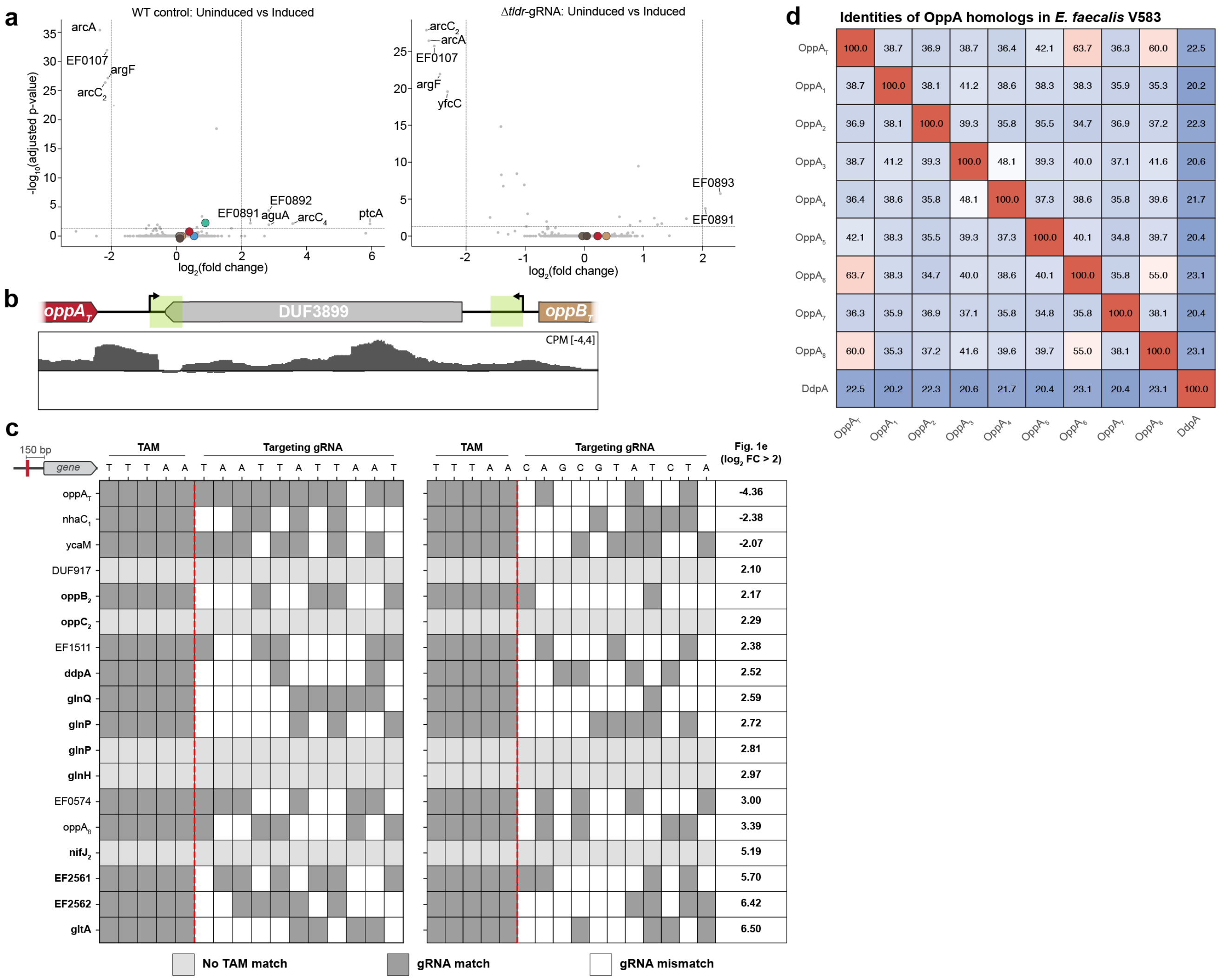


**Figure S2 | Controls supporting the specificity of TldR-mediated *oppA*$_T$ repression in *E. faecalis*. a,** Differential expression between uninduced and induced conditions for the WT control strain (left) and the Δ*tldr*-gRNA strain (right). Volcano plots show $\log_2$ fold change (x-axis) versus adjusted p-value (y-axis). Dashed lines indicate significance thresholds ($|\log_2 FC| > 2$, $p_{adj} < 0.05$). Highlighted genes of the *opp*$_T$ operon are colored as in **Figure 1** and remain near baseline in both strains, indicating no agmatine-dependent effect on this operon. Labeled genes (e.g., *arcA*, *arcC*$_2$, *arcC*$_4$, *argF*, *aguA*) encode arginine/agmatine catabolic enzymes that respond directly to agmatine induction, serving as a positive control for successful induction. **b,** Representative RNA-seq coverage track of the intergenic region between *oppA*$_T$ and *oppB*$_T$ in the WT control strain. Promotech-predicted[69] promoter regions within the track are highlighted in green. The coverage dip between the two predicted promoters is consistent with *oppA*$_T$ being transcribed independently of the downstream *oppB–F*$_T$*–tldR* operon. **c**, Best sequence match for a contiguous TAM-gRNA sequence (targeting, left; non-targeting, right) found within a 150-bp window upstream of the start codon for all genes significantly differentially expressed (|log2FC| > 2, padj < 0.05) in **Figure 1e**. **d,** Pairwise amino acid sequence identity matrix for OppA homologs encoded by *E. faecalis* V583, including the TldR-associated OppA$_T$, eight additional OppA paralogs, and the related substrate-binding protein DdpA. Percent identity values are indicated within each cell and color-coded. These data highlight the diversity of substrate-binding proteins coexisting within a single genome, consistent with functional specialization.

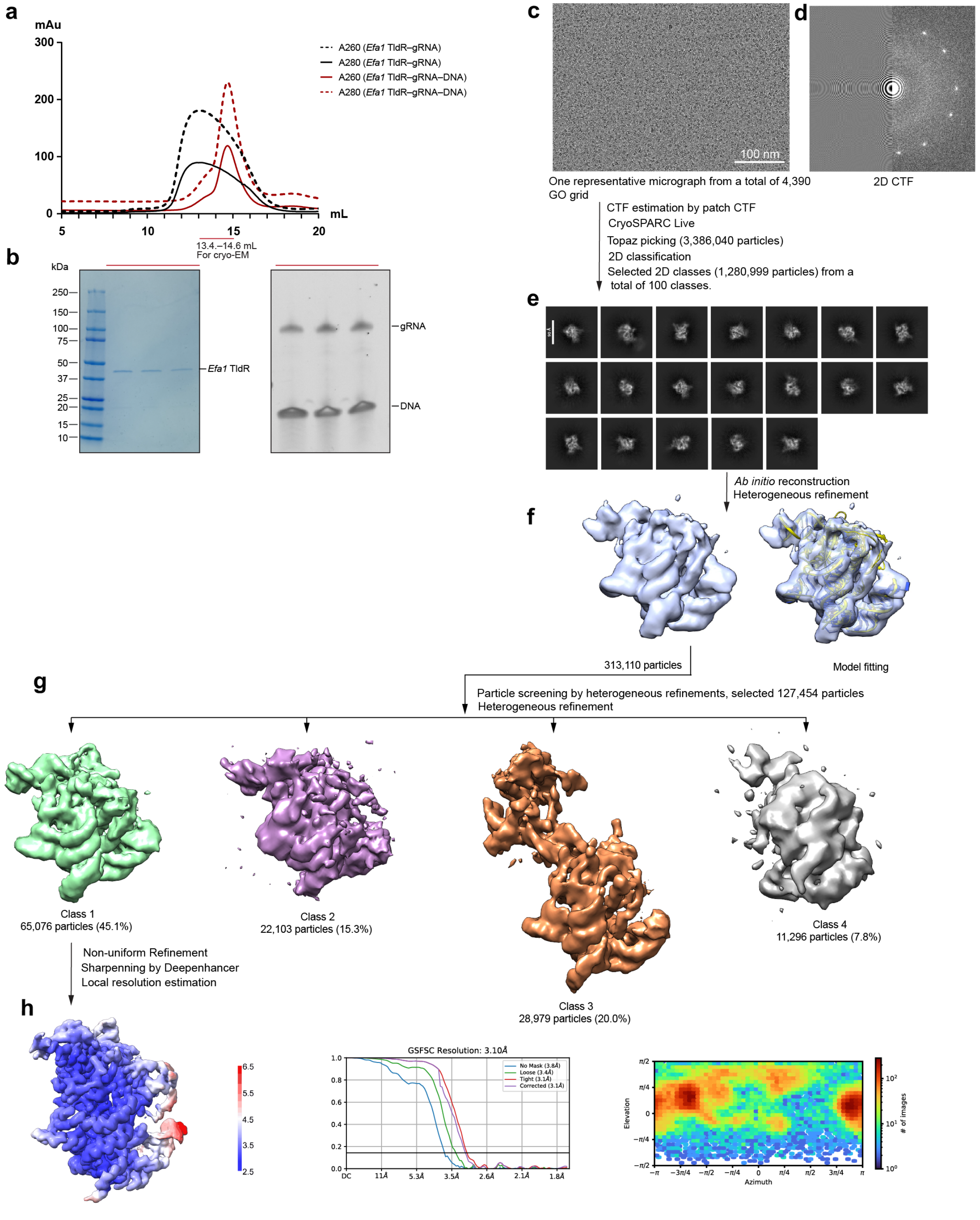


**Figure S3 | Sample preparation and cryo-EM analysis of *Efa$_1$*TldR–gRNA–DNA ternary complex. a,** Size exclusion chromatography (SEC) profiles of *Efa$_1$*TldR–gRNA (black) and the *Efa$_1$*TldR–gRNA–DNA complex (red). UV absorbance curves at 280 nm and 260 nm are shown in solid and dashed lines, respectively. **b,** Corresponding SDS-PAGE (left) and urea-PAGE (right) analyses from the SEC experiment in **a**. **c,** Representative cryo-EM micrograph from a total of 4,390 micrographs. **d,** 2D CTF estimation of the representative micrograph in **c**. **e,** Representative 2D class averages. **f,** First round of heterogeneous refinement with four classes; only the best class is shown. **g,** Four classes from the second round of heterogeneous refinement. **h,** Local resolution map of the final non-uniform refinement of *Efa1*TldR–gRNA–DNA ternary complex at 3.10 Å; global gold-standard FSC curves and particle angular distributions are shown on the right.

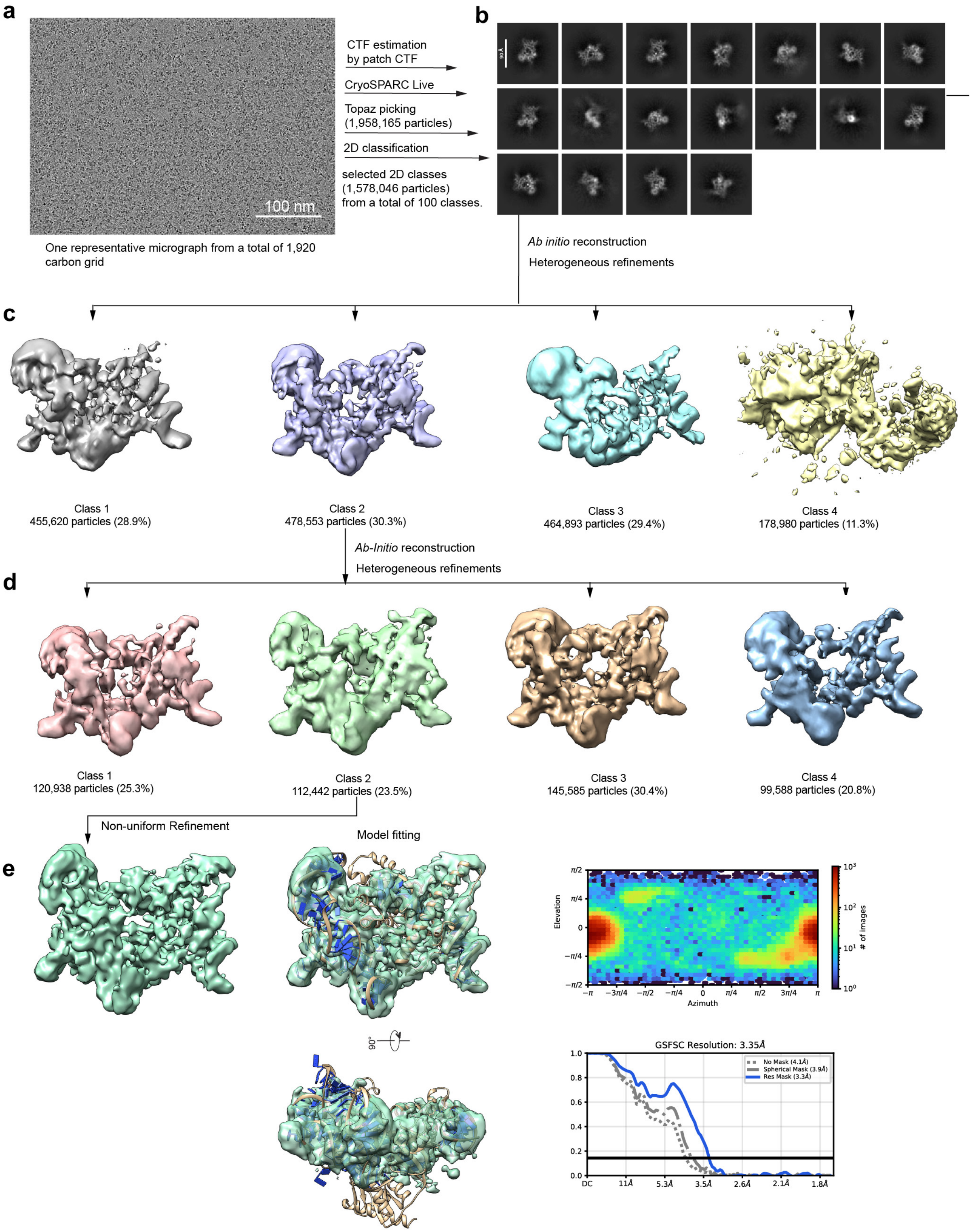


**Figure S4 | Cryo-EM analysis of the *Efa$_1$*TldR–gRNA–DNA ternary complex on carbon grids. a,** A representative cryo-EM micrograph from a total of 1,920 micrographs. **b,** Representative 2D class averages. **c,** First round of heterogeneous refinement with four classes. **d,** Second round of heterogeneous refinement of the 478,553 particles in Class 2 from **c**, again yielding four classes. **e,** Final non-uniform refinement of *Efa$_1$*TldR–gRNA–DNA complex at 3.35 Å. The reconstruction shows incomplete and anisotropic density consistent with preferred particle orientation; global gold-standard FSC curves and particle angular distributions are shown on the right.

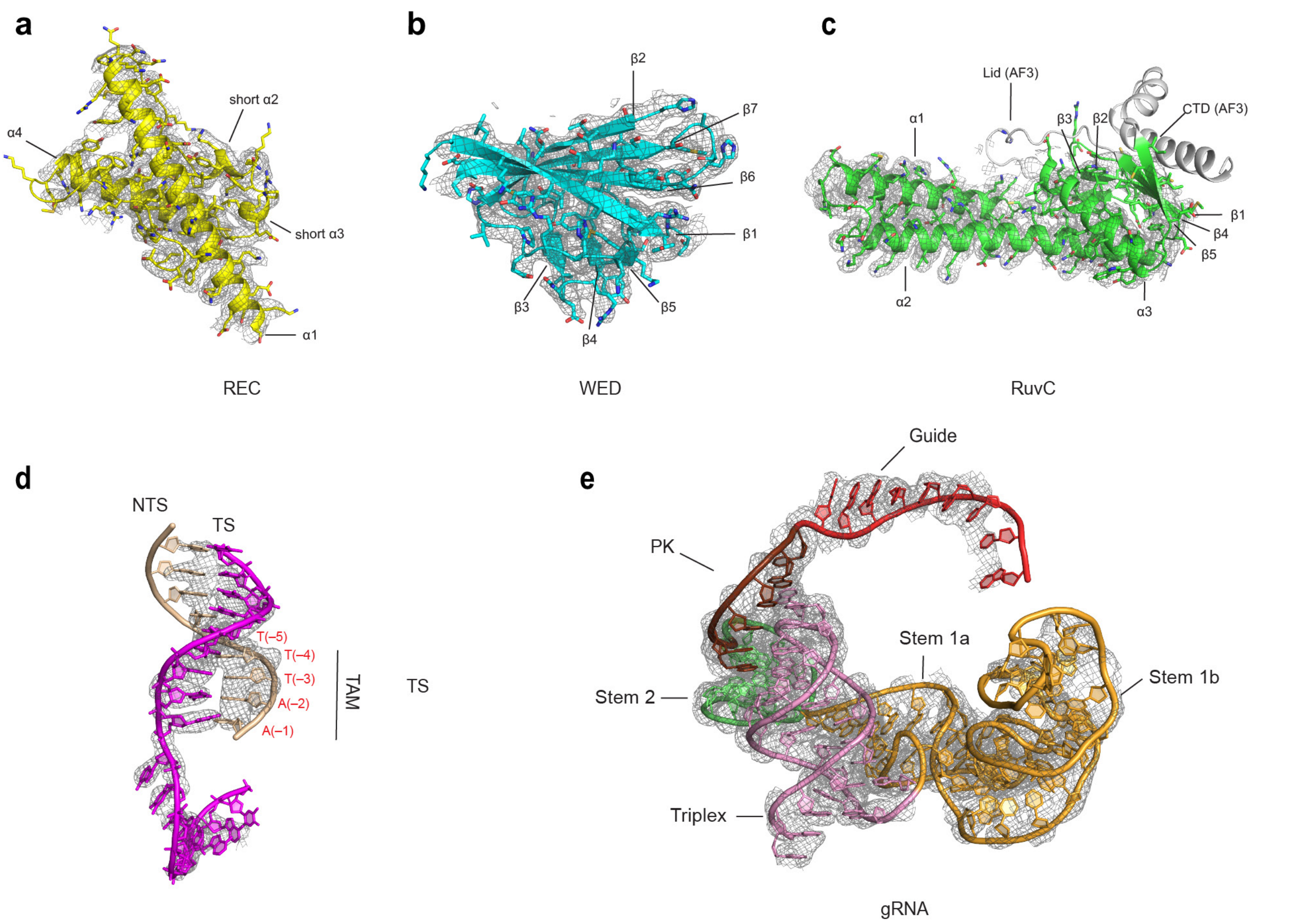


**Figure S5 | Cryo-EM density and atomic models of the *Efa$_1$*TldR–gRNA–DNA ternary complex. a–c,** Cryo-EM density map (mesh) with fitted atomic models for the REC (**a**), WED (**b**), and RuvC (**c**) domains. Regions corresponding to the Lid motif and CTD were not resolved and are shown as predicted AlphaFold 3 models. **d,** Cryo-EM density and model of the target DNA duplex, with the TAM region indicated. **e,** Cryo-EM density and model of the gRNA, with structural elements annotated.

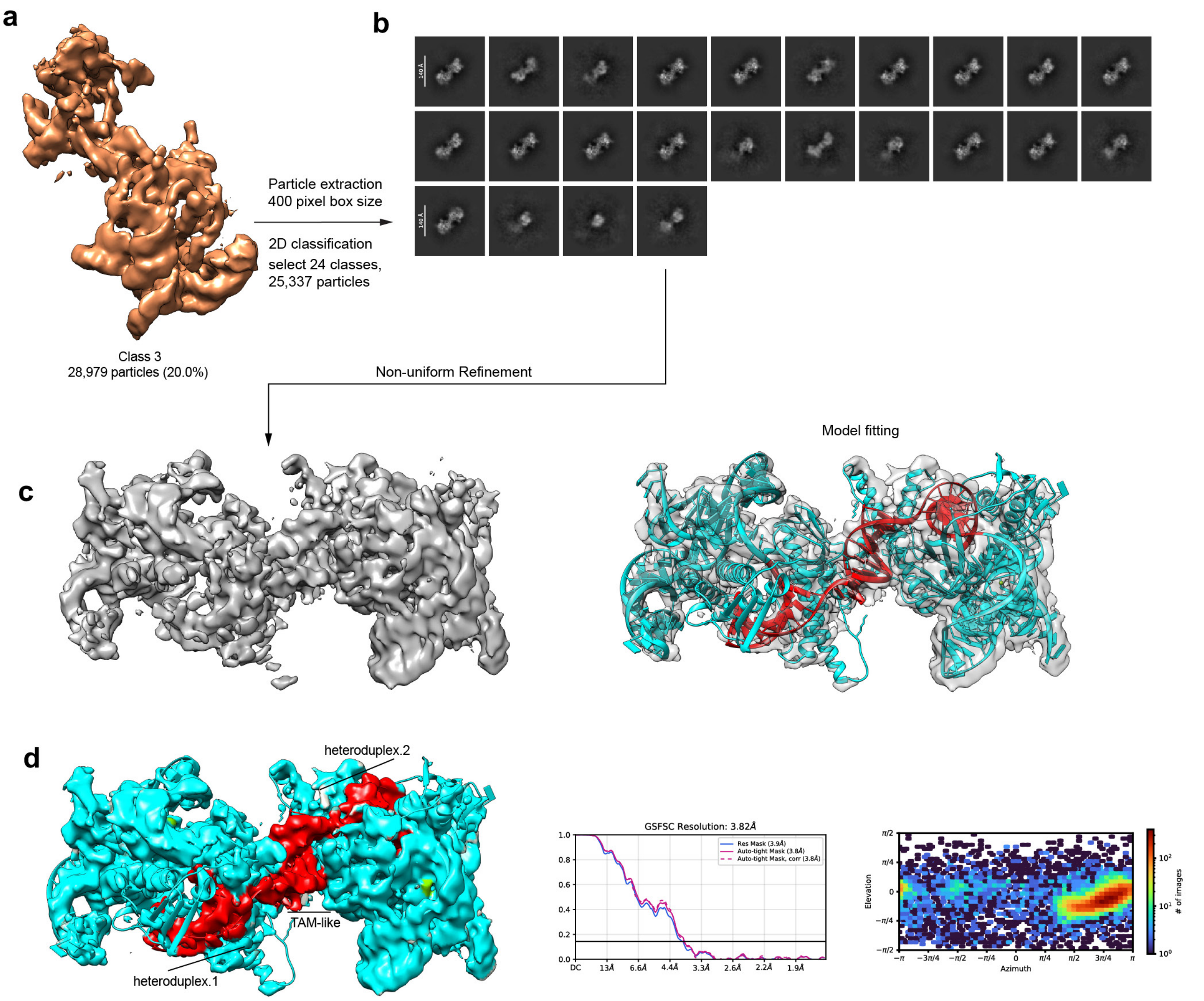


**Figure S6 | Cryo-EM analysis of a head-to-head TldR ternary dimer. a,** Cryo-EM density map of the dimeric assembly corresponding to the class shown in **Fig. S3g**. **b,** Representative 2D class averages following particle re-extraction with an enlarged box size. **c,** Non-uniform refinement yielded a 3.82 Å reconstruction, with the fitted atomic model shown at right. **d,** Surface representation highlighting the RNA–DNA heteroduplex within each protomer and a single TAM-like DNA duplex (red) positioned at the dimer interface. The head-to-head arrangement may arise from non-canonical base pairing between the two target DNA strands. Global gold-standard FSC curves and particle angular distributions are shown at right.

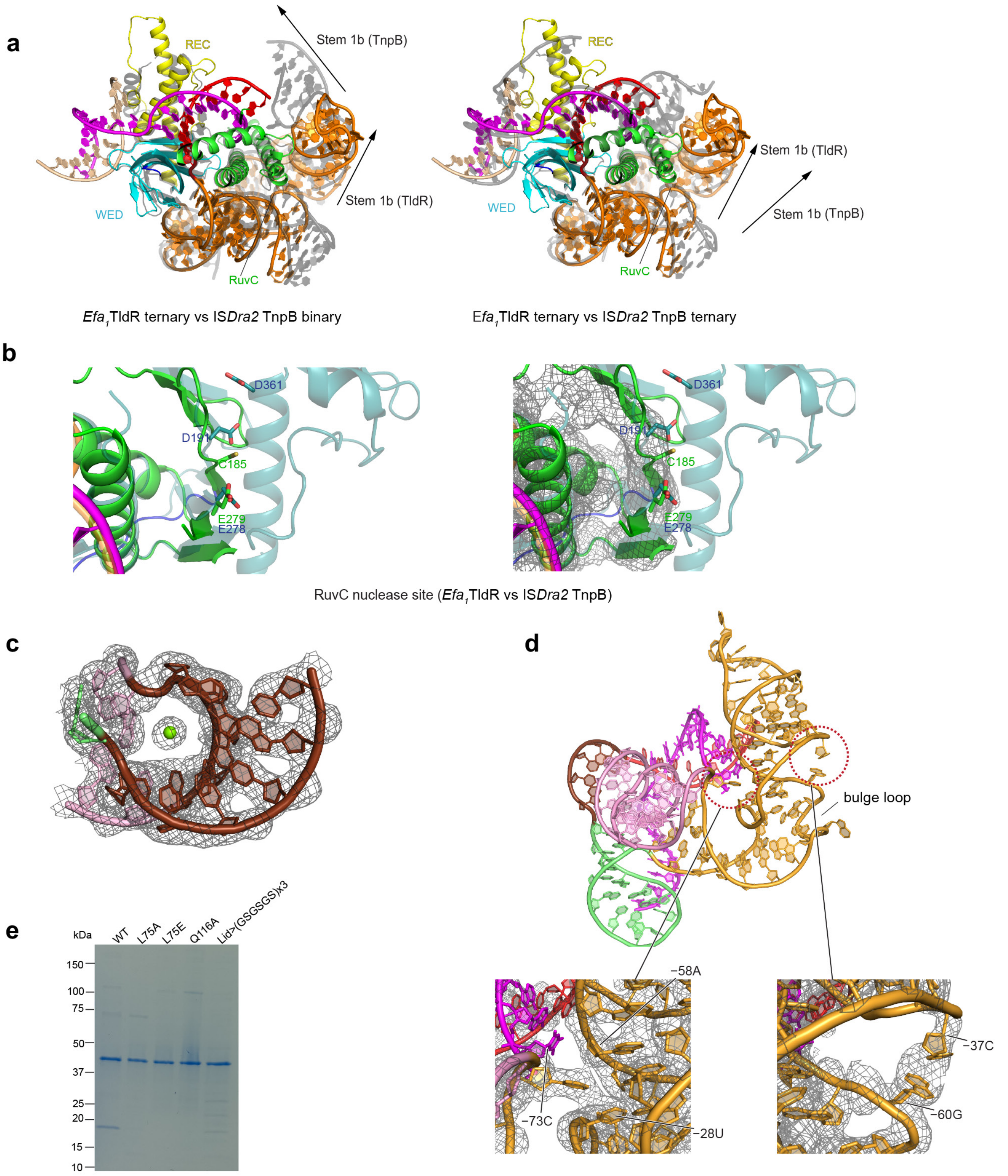


**Figure S7 | Structural comparison of *Efa₁*TldR and IS*Dra2* TnpB and unique features of the TldR gRNA. a,** Superimposition of the *Efa*$_1$TldR ternary complex with the IS*Dra2* TnpB binary complex (left) and ternary complex (right), shown in grey. **b,** Comparison of the RuvC nuclease active site from TldR (green) and TnpB (blue), highlighting substitution of catalytic residues in TldR. Cryo-EM density for the TldR RuvC domain is shown in mesh at right. **c,** Vertical view of the gRNA pseudoknot (PK) region, with the coordinated $Mg^{2+}$ ion indicated (green sphere). **d,** Side view of *Efa*$_1$TldR stem 1b, with magnified insets below highlighting select regions. **e**, SDS–PAGE analysis of purified wild-type and mutant TldR–gRNA complexes.

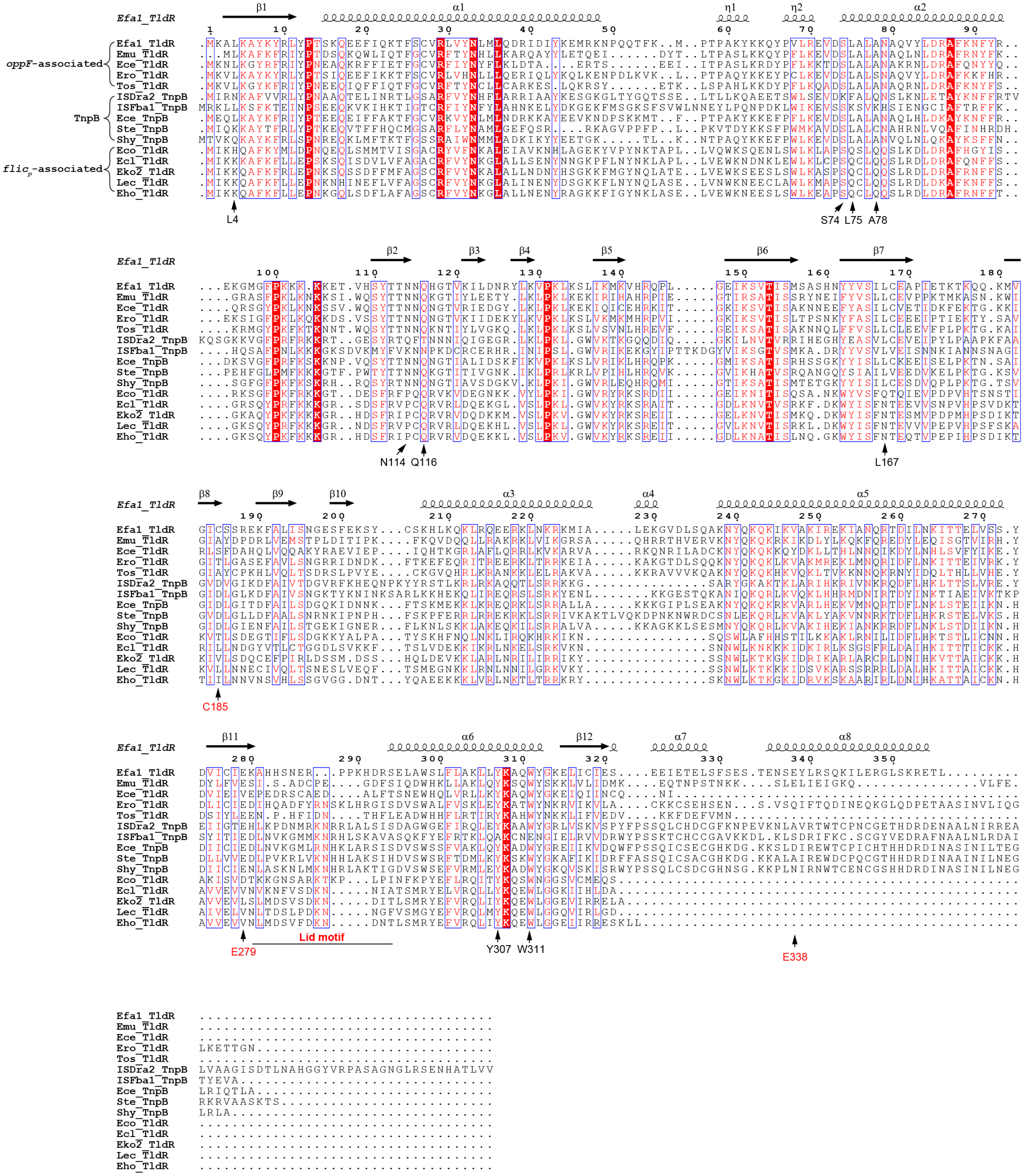


**Figure S8 | Sequence alignment of representative *oppF*-associated and *fliC*-associated TldRs and TnpB homologs.** Multiple sequence alignment with secondary structure elements of $Efa_1$TldR indicated above. Conserved residues are highlighted, and positions involved in TAM recognition and heteroduplex stabilization are annotated. The Lid motif and regions corresponding to the RuvC catalytic site are indicated in red.

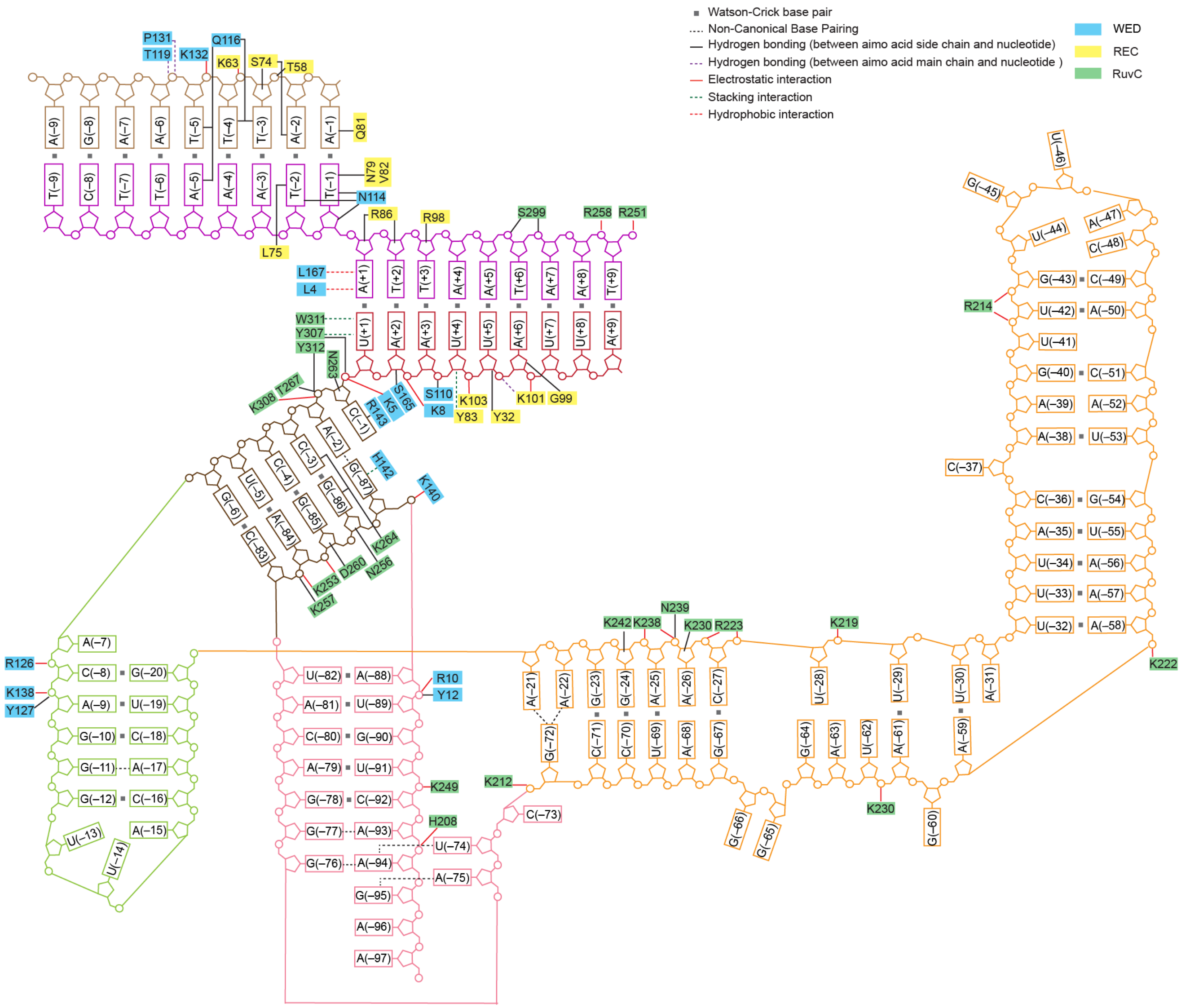


**Figure S9 | Schematic of gRNA–DNA interactions within the *Efa$_1$*TldR ternary complex.** Schematic representation of the gRNA and target DNA, with protein–nucleic acid interactions annotated; residues are color-coded by domain: REC (yellow), WED (blue), and RuvC (green). Interaction types — including Watson-Crick and non-canonical base pairing, side-chain and main-chain hydrogen bonding, electrostatic interactions, stacking, and hydrophobic contacts — are indicated by the colored and dashed lines defined in the key.

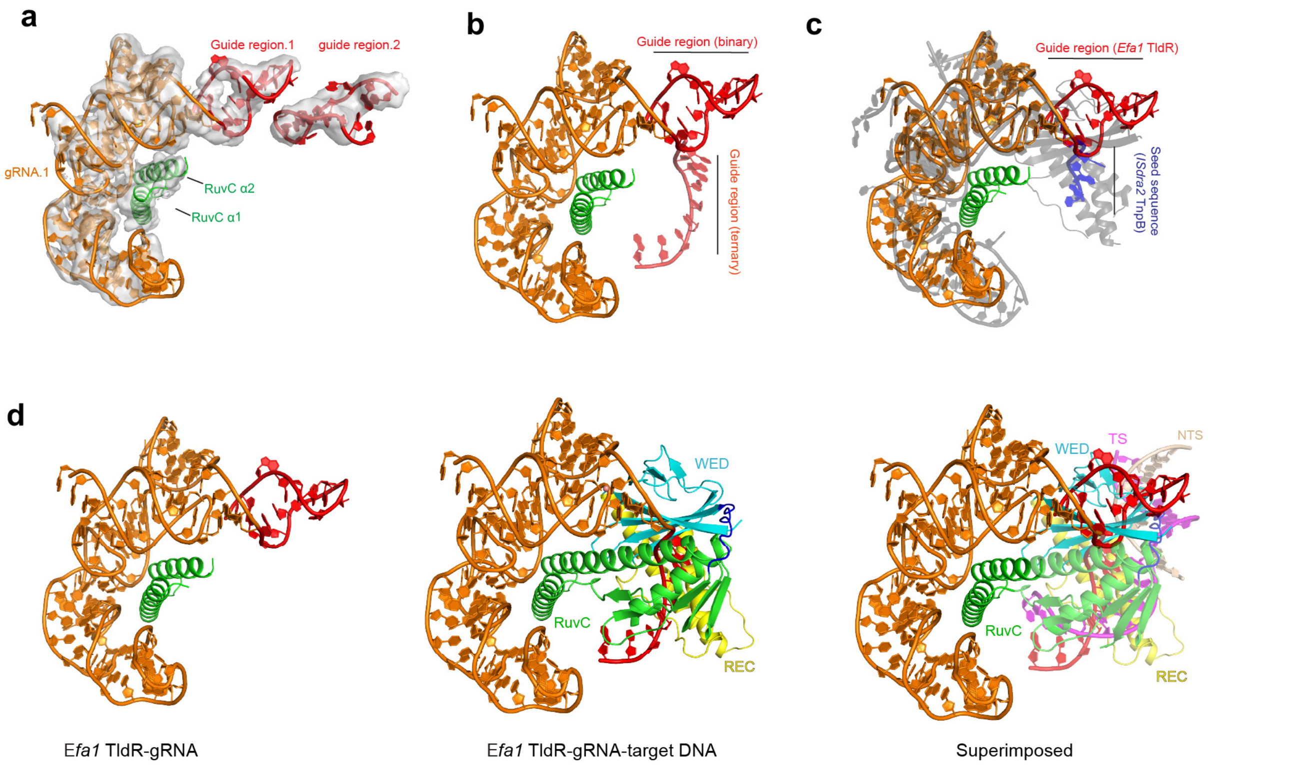


**Figure S10 | Overall structure of *Efa*$_1$TldR–gRNA binary complex. a,** Cryo-EM density map with fitted atomic model. **b,** Comparison of the guide region between the binary and ternary complexes. **c,** Superimposition of the *Efa*$_1$TldR binary complex with the IS*Dra2* TnpB binary complex (grey). **d,** Superimposition of the *Efa*$_1$TldR binary and ternary complexes, showing that the guide region of the binary complex would clash with the WED domain in the ternary complex.

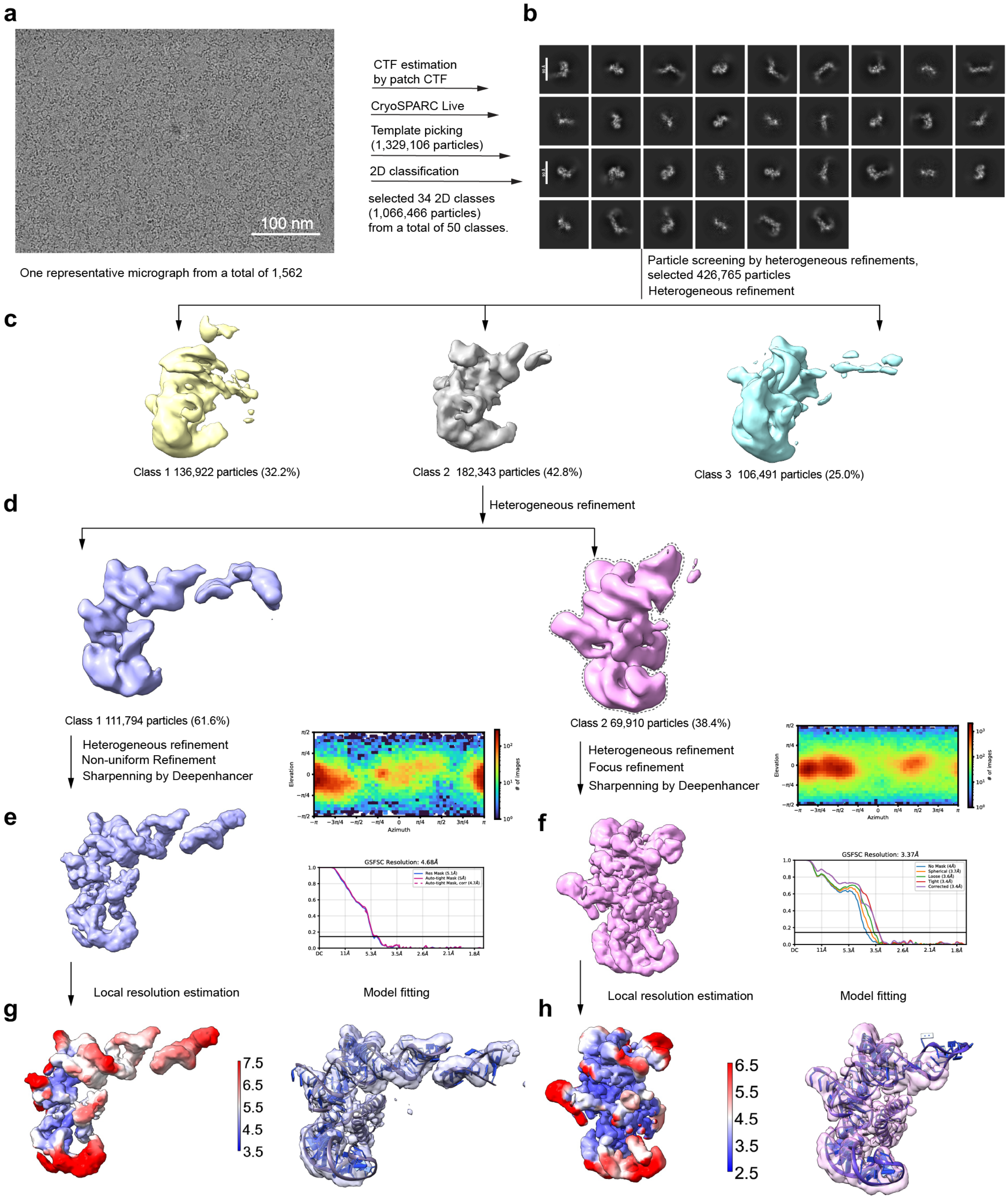
a
100 nm
One representative micrograph from a total of 1,562
CTF estimation
by patch CTF
CryoSPARC Live
Template picking
(1,329,106 particles)
2D classification
selected 34 2D classes
(1,066,466 particles)
from a total of 50 classes.
b
Particle screening by heterogeneous refinements,
selected 426,765 particles
Heterogeneous refinement
c
Class 1 136,922 particles (32.2%)
Class 2 182,343 particles (42.8%)
Class 3 106,491 particles (25.0%)
d
Heterogeneous refinement
Class 1 111,794 particles (61.6%)
Class 2 69,910 particles (38.4%)
Heterogeneous refinement
Non-uniform Refinement
Sharpenning by Deepenhancer
Heterogeneous refinement
Focus refinement
Sharpenning by Deepenhancer
e
f
Elevation
Azimuth
### of images
GSFSC Resolution: 4.68Å
GSFSC Resolution: 3.37Å
Local resolution estimation
Model fitting
Local resolution estimation
Model fitting
g
h
7.5
6.5
5.5
4.5
3.5
6.5
5.5
4.5
3.5
2.5

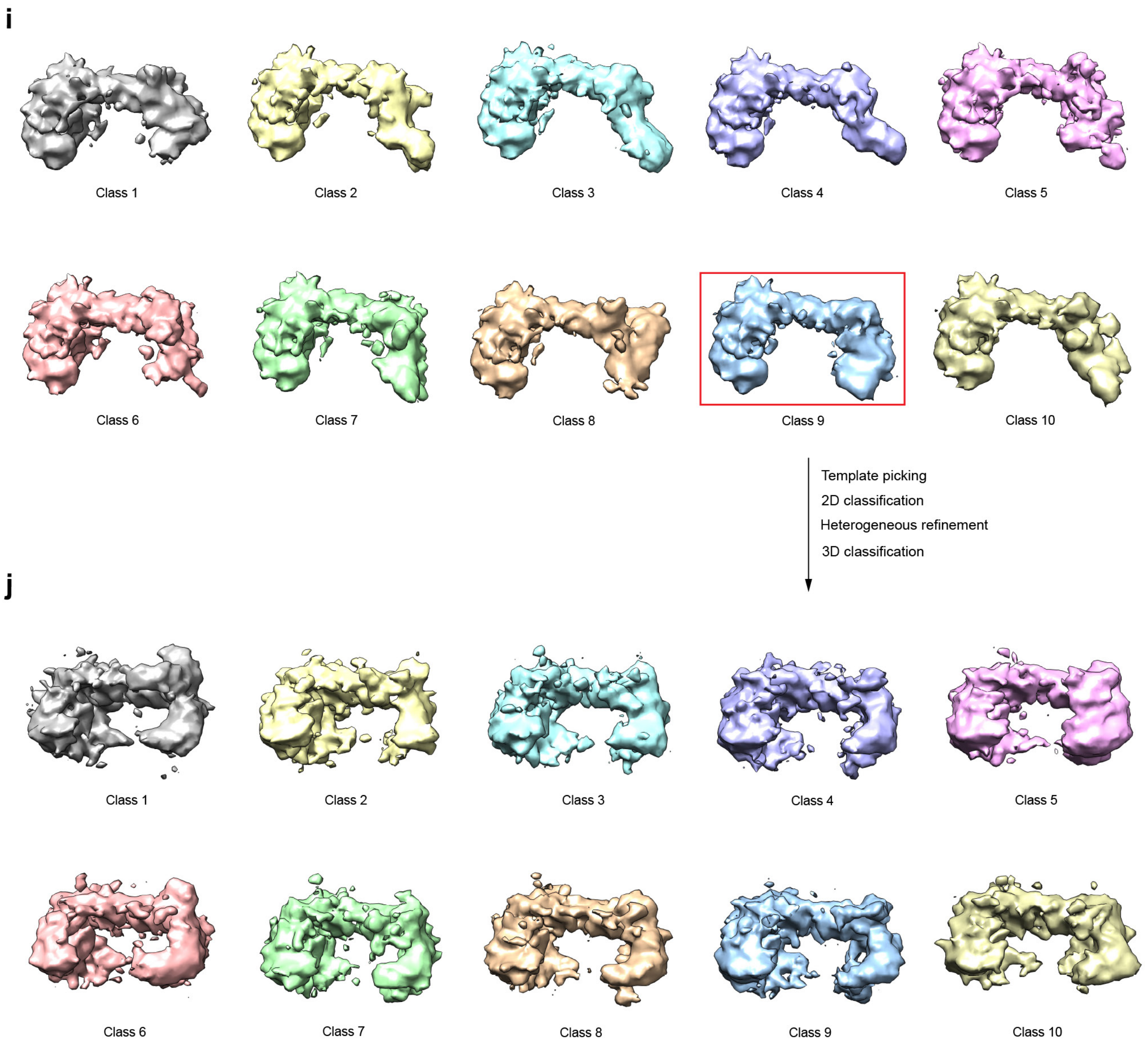


**Figure S11 | Cryo-EM data processing of the *Efa*$_1$TldR–gRNA binary complex. a,** Representative cryo-EM micrograph from a total of 1,562 micrographs. **b,** Representative 2D class averages. **c,** Initial particle classes following heterogeneous refinement. **d,** Second round of heterogeneous refinement of the 182,343 particles in Class 2 from **c**, yielding two classes. **e,** Final cryo-EM reconstruction of the dimeric assembly at 4.68 Å; global gold-standard FSC curves and particle angular distributions are shown on the right. **f,** Final focused refinement of the *Efa*$_1$TldR–gRNA single protomer at 3.37 Å; global gold-standard FSC curves and particle angular distributions are shown on the right. **g,h,** Local resolution maps and fitted atomic models for the dimeric assembly (**g**) and single protomer (**h**). **i,** Ten representative classes from 3D classification using particles from **g**. **j**, Refined classes following re-processing of the boxed class (Class 8) from *i* through template picking, 2D classification, heterogeneous refinement, and a second round of 3D classification.

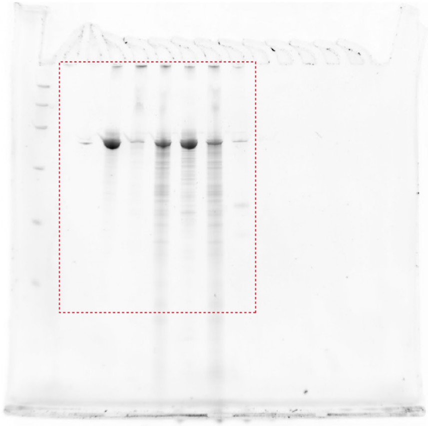

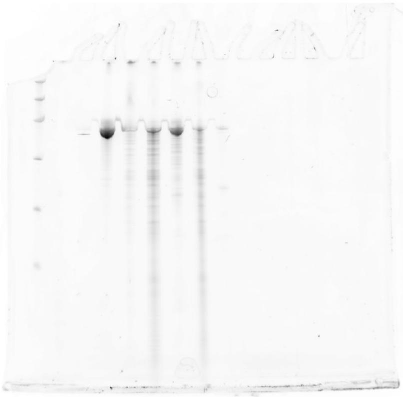

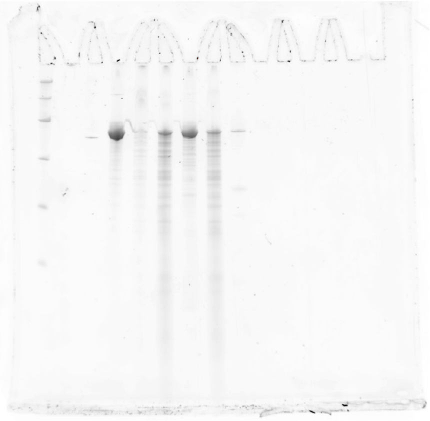

**Figure 4d**
repeat 1

**Figure 4d**
repeat 2

**Figure 4d**
repeat 3

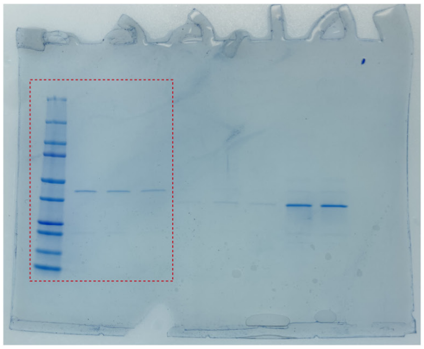

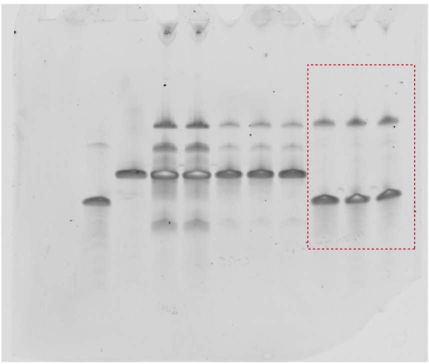

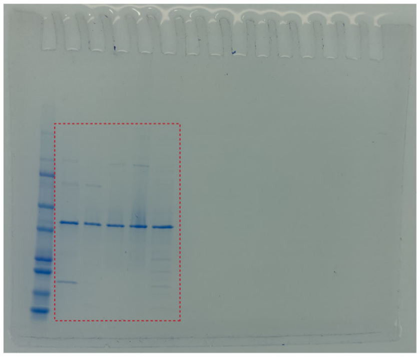

**Figure S3b**
*Efa*$_1$TldR

**Figure S3b**
gRNA and target DNA

**Figure S7e**
*Efa*$_1$TldR mutants

**Figure S12 | Source data for gel-based assays.** Uncropped and unprocessed gel images corresponding to experimental data shown in **Fig. 4d** (top row, three biological replicates) and **Fig. S3b** (bottom row, left and center: SDS-PAGE and urea-PAGE of *Efa*$_1$TldR–gRNA and target DNA) and **Fig. S7e** (bottom row, right: SDS-PAGE of *Efa*$_1$TldR mutants). Regions corresponding to the cropped/processed images shown in the main and supplementary figures are indicated by dashed boxes.
